# Rapid reversible osmoregulation of cytoplasmic biomolecular condensates of human interferon-α-induced antiviral MxA GTPase

**DOI:** 10.1101/2022.08.30.505854

**Authors:** Pravin B. Sehgal, Huijuan Yuan, Ye Jin

**Affiliations:** Departments of Cell Biology and Anatomy, New York Medical College, Valhalla, New York, USA; Departments of Medicine, New York Medical College, Valhalla, New York, USA

**Author notes:** For correspondence: Pravin B. Sehgal. Huijuan Yuan: Division of Pulmonary, Allergy, and Critical Care Medicine, Department of Medicine, University of Pittsburgh School of Medicine, Pittsburgh, Pennsylvania, USA. Ye Jin: Department of Immunology, University of Pittsburgh School of Medicine, Pittsburgh, Pennsylvania, USA. Contact information: Dr. Pravin B. Sehgal, Professor, Dept. Cell Biology and Anatomy, New York Medical College, Rm. A17 Basic Science Building, Valhalla, NY 10595.

## Abstract

Cellular and tissue-level edema is a common feature of acute viral infections such as covid-19, and of many hyponatremic hypoosmolar disorders. However, there is little understanding of the effects of cellular edema on antiviral effector mechanisms. We previously discovered that cytoplasmic human MxA, a major antiviral effector of Type I and III interferons against several RNA- and DNA-containing viruses, existed in the cytoplasm in phase-separated membraneless biomolecular condensates of varying sizes and shapes. In this study we investigated how hypoosmolar conditions, mimicking cellular edema, might affect the structure and antiviral function of MxA condensates. Cytoplasmic condensates of both IFN-α-induced endogenous MxA and of exogenously expressed GFP-MxA in human A549 lung and Huh7 hepatoma cells rapidly disassembled within 1-2 min when cells were exposed to hypotonic buffer (∼ 40-50 mOsm), and rapidly reassembled into new structures within 1-2 min of shifting of cells to isotonic culture medium (∼ 330 mOsm). MxA condensates in cells continuously exposed to culture medium of moderate hypotonicity (in the range one-fourth, one-third or one-half isotonicity; range 90-175 mOsm) first rapidly disassembled within 1-3 min, and then, in most cells, spontaneously reassembled 7-15 min later into new structures. Condensate reassembly, whether induced by isotonic medium or occurring spontaneously under continued moderate hypotonicity, was preceded by “crowding” of the cytosolic space by large vacuole-like dilations (VLDs) derived from internalized plasma membrane. Remarkably, the antiviral activity of GFP-MxA against vesicular stomatitis virus survived hypoosmolar disassembly. Overall, the data highlight the exquisite sensitivity of MxA condensates to rapid reversible osmoregulation.

## Introduction

Hyponatremia is the commonest electrolyte disturbance, and is a clinical feature in 15-20% of emergency room admissions (1–6). Hypotonic (hypoosmolar) hyponatremia is the largest underlying cause of this disorder in situations such as salt-wasting nephropathies (e.g. polycystic kidney disease), mineralocorticoid deficiency (e.g. Addison’s disease), cirrhosis, congestive heart disease, cancer and other situations characterized by tissue-level edema-formation (3, 5, 6). Overall, general morbidity and mortality is higher in such patients across a wide range of underlying conditions, with neurologic symptoms a common manifestation (1–6). At the cellular level complex biochemical and membrane trafficking mechanisms come into play in order to maintain intracellular homeostasis (5, 6, 7–14). These include stimulation of endocytosis leading to rapid plasma membrane internalization by swollen cells into vacuole-like dilations (VLDs)(7, 9, 12, 13), activation of transient mechanosensitive receptor ion channels, activation of receptor tyrosine kinases, relocalization of Src in membrane blebs, and F-actin diasassembly including inhibition of myosin light chain kinase in response to cellular “osmosensing” (7–14). Hyperosmotic stress (HOPS) also triggers cellular homeostatic mechanisms including phase separation and condensate formation by multivalent target proteins such as the P-body trimeric protein DCP1A (15–17).

Tissue-level and cellular edema is a common feature of inflammation, especially following cytopathic virus infection (18–22). Already in 1966, it was remarked that following vaccinia virus infection “the most prominent early cytopathic change was intracellular edema evidenced by low electron density to the background cytoplasmic material and dilatation of endoplasmic reticulum” (22). More recently, the pathogenesis of the pulmonary edema observed in patients with Covid-19 has attracted attention (23–25). While Type I and III interferons have been shown to inhibit SARS-CoV-2 replication in cell culture, and defects in the IFN system (auto-antibodies and receptor mutations) have been implicated in severe/fatal Covid-19 disease (18, 26–31), there is little understanding whether IFNs preserve their antiviral activity against SARS-CoV-2 or other target viruses in edematous cells i.e. in cells under hypotonic conditions. More generally, how do antiviral defenses function in hypoosmolar disorders?

It is now increasingly recognized that liquid-liquid phase-separation (LLPS) leads to the formation of biomolecular condensates [also called membraneless organelles (MLOs)] in the cytoplasm and nucleus of eukaryotic cells (32–39). MLOs provide a scaffold for diverse cellular functions (e.g. cell signaling, nuclear transcription, RNA splicing and processing, mRNA storage and translation, DNA sensing, synaptic function, and mitosis)(32–39). It is also now recognized that the replication of many viruses in mammalian cells involves LLPS condensates [e.g. in the life cycles of vesicular stomatitis (VSV), rabies (Negri bodies), influenza A, respiratory syncytial, Ebola, measles, Epstein-Barr, and SARS-CoV-2 viruses](reviewed in 38, 39).

Four years ago, we made the discovery that human myxovirus resistance protein MxA, a 62-kDa dynamin-family large GTPase which is induced 100-300-fold in diverse cell-types by Type I IFNs (α/β families) and Type III IFNs (λ family)(40–42), formed phase-separated membraneless organelles (MLOs) in the cytoplasm (43, 44). MxA is a major antiviral effector of IFNs-α/β and IFN-λ against diverse RNA- and DNA-viruses (40–42). Cytoplasmic MxA condensates, of gel-like internal consistency, associated with the viral nucleocapsid (N) protein of a target viruses [e.g. vesicular stomatitis virus (VSV) in cells exhibiting an antiviral phenotype.(44); also see Kochs et al 2002 for very early data for La Crosse virus N protein in membraneless MxA structures (45) by electron microscopy]. Remarkably, condensates formed by *exogenously* expressed GFP-MxA in human Huh7 and Hep3B hepatoma cells showed rapid disassembly (within 1-2 min) in cells exposed to hypotonic medium, and rapid reassembly (within 1-2 min) into new condensate structures when cells were subsequently shifted to isotonic medium (44, 46). In the present study we investigated whether IFN-α-induced *endogenous* MxA structures in lung-derived cells showed osmosensing properties, the underlying mechanisms, and whether osmotic disassembly/reassembly of MxA condensates affected their antiviral activity (against VSV). The data obtained highlight the rapid hypotonicity-driven disassembly and, subsequent, isotonicity-driven or even *spontaneous* reassembly of MxA condensates in lung cells. Remarkably, the antiviral activity of MxA survived osmotic cycling. The observation that cell integrity was required for maintenance of MxA higher-order structures in the cytoplasm places limitations on interpretation of prior data on MxA oligomerization derived from previous solution-based analyses (40–42). More generally, the present observations point to similar limitations in biochemical studies in free solution of other proteins which form higher-order clusters and condensates in the cytosol in intact cells (e.g. the transcription factor STAT3; 47, 48).

## Results

### Properties of endogenous MxA condensates/granules in human lung cells

Our previous studies of cytoplasmic MxA condensates were largely carried out using GFP-tagged human MxA transiently expressed in human hepatoma (Huh7 and Hep3B) cells (44). These GFP-MxA condensates were variably sized and shaped, showed homotypic fusion, were disassembled by 1,6-hexanediol, were membraneless by thin-section EM in a correlated light and electron microscopy (CLEM) assay, and had an internal gel-like consistency by fluorescence recovery after photobleaching (FRAP) assay (44). Cells expressing GFP-MxA condensates showed an antiviral phenotype against VSV; in many such cells the VSV N protein associated with GFP-MxA condensates (44). Unexpectedly, we observed that GFP-MxA condensates disassembled in 1-2 min in cells exposed to hypotonic medium (ELB; erythrocyte lysis buffer; 40-50 mOsm), and reassembled into new structures within 1-2 min of shifting cells to isotonic culture medium or phosphate-buffered saline (PBS)(∼330 mOsm). Even in isotonic medium, the integrity of GFP-MxA condensate structures required an intact plasma membrane in that the addition of saponin (0.03%) to the culture medium caused rapid disassembly of the bulk of GFP-MxA condensates, leaving behind a saponin-resistant core in 10-15% of the transfected cells (44). With increasing evidence of the involvement of IFN-α as a protective mechanism in covid-19 caused by the SARS-CoV-2 virus, the common development of pulmonary edema in such patients, and observations showing increased levels of MxA (often referred to as Mx1 in clinical papers), in lung tissues from covid-19 patients (18–31), we investigated MxA structures formed by IFN-α-treated human lung-derived cells (the A549 adenocarcinoma line; and the SARS-CoV-2 permissive hACE2-A549 cells) under isotonic and hypoosmolar conditions.

The Western blot data in Fig. 1A confirm that exposure of A549 human lung-derived cells in culture to IFN-α2 enhanced expression of cellular MxA by more than one-hundred fold. By immunofluoresnece analyses this endogenous MxA protein was cytoplasmic, and was in metamorphic granular structures (Fig. 1B). We used the method of Fourier transformation-based “minimum” size-filter of objects using Image J software to estimate the amount of MxA in condensates vs diffuse in the cytosol (explained in Fig. 2 in Sehgal et al, ref. 38; illustrated in Fig. 7B below). Overall, approximately 60-70% of endogenous MxA in IFN-α-treated A549 cells was estimated to be in condensed/granular structures (Fig. 1B). This is an estimate similar to that for exogenously expressed GFP-MxA (Figs. 3A, 6 and 7).

**Figure 1.**
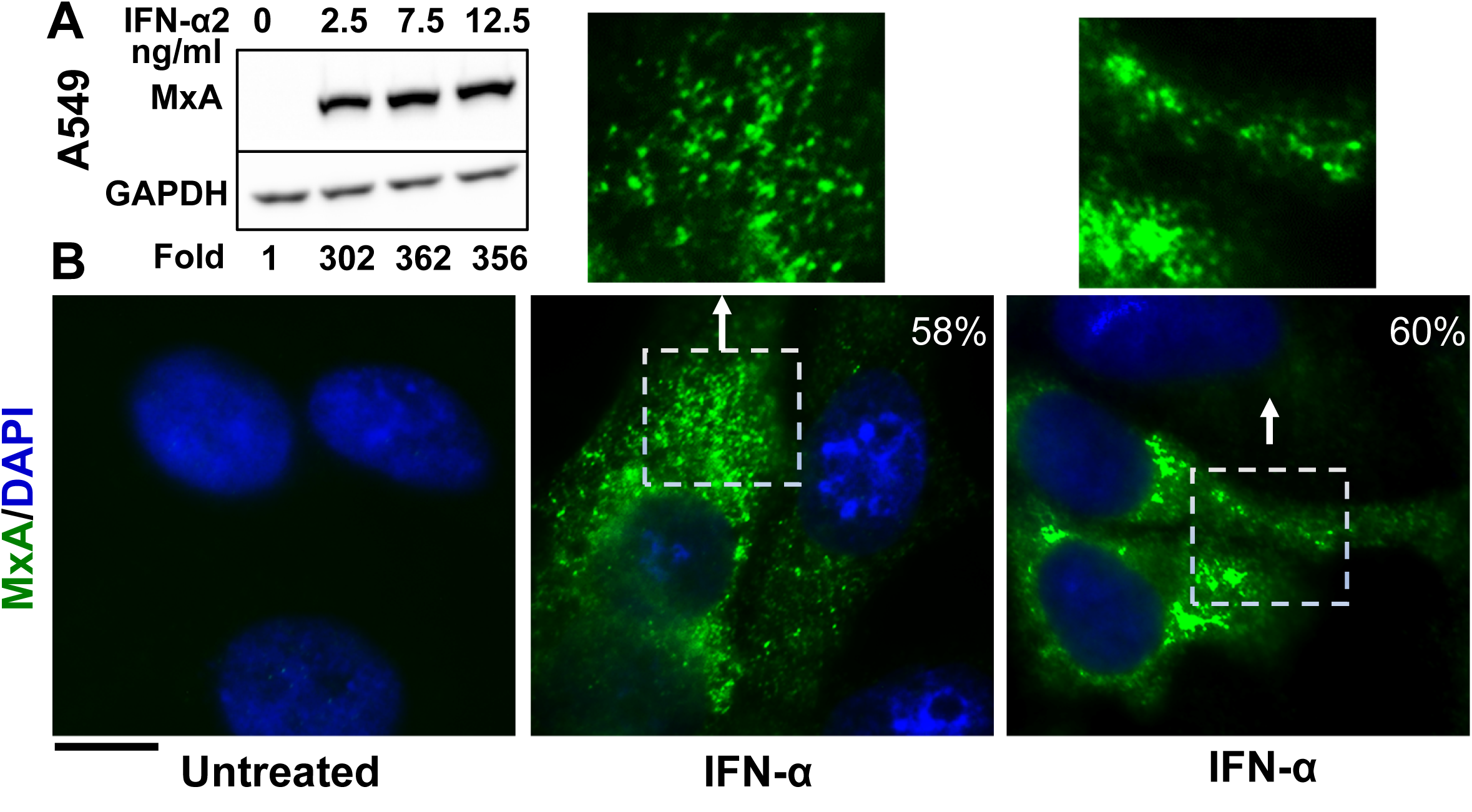
Cytoplasmic granules of endogenous MxA induced by IFN-*α*2 in human lung-derived A549 cells. Panel A, A549 cells in culture (in 35 mm plates) were exposed to IFN-α2 at the indicated concentrations for 2 days, washed with PBS, and whole-cell extracts prepared using buffer isotonic buffer containing 0.1% SDS and 0.5% Triton X-100 (47, 57). Matching protein aliquots (30 µg) were Western blotted for MxA and GAPDH, and fold-induction of MxA by IFN evaluated using Image J. Panel B. A549 cells in 35 mm plates without or with exposure to IFN-α2 (20 ng/ml) for 2 days, were fixed (4% PFA in PBS for 1 hr at 4°C), permeabilized (using digitonin or saponin buffer; 57), immunostained for MxA (scale bar = 10 µm). The boxed insets are also immustrated at higher magnification. Quantitation of MxA in respective images present in condensates as a percent of the total is indicated in respective panels; this was evaluated using a Fourier filter in Image J to subtract objects up to and below a designated pixel radius (discussed in ref. 38; see Fig. 7B for an example).

**Figure 2.**
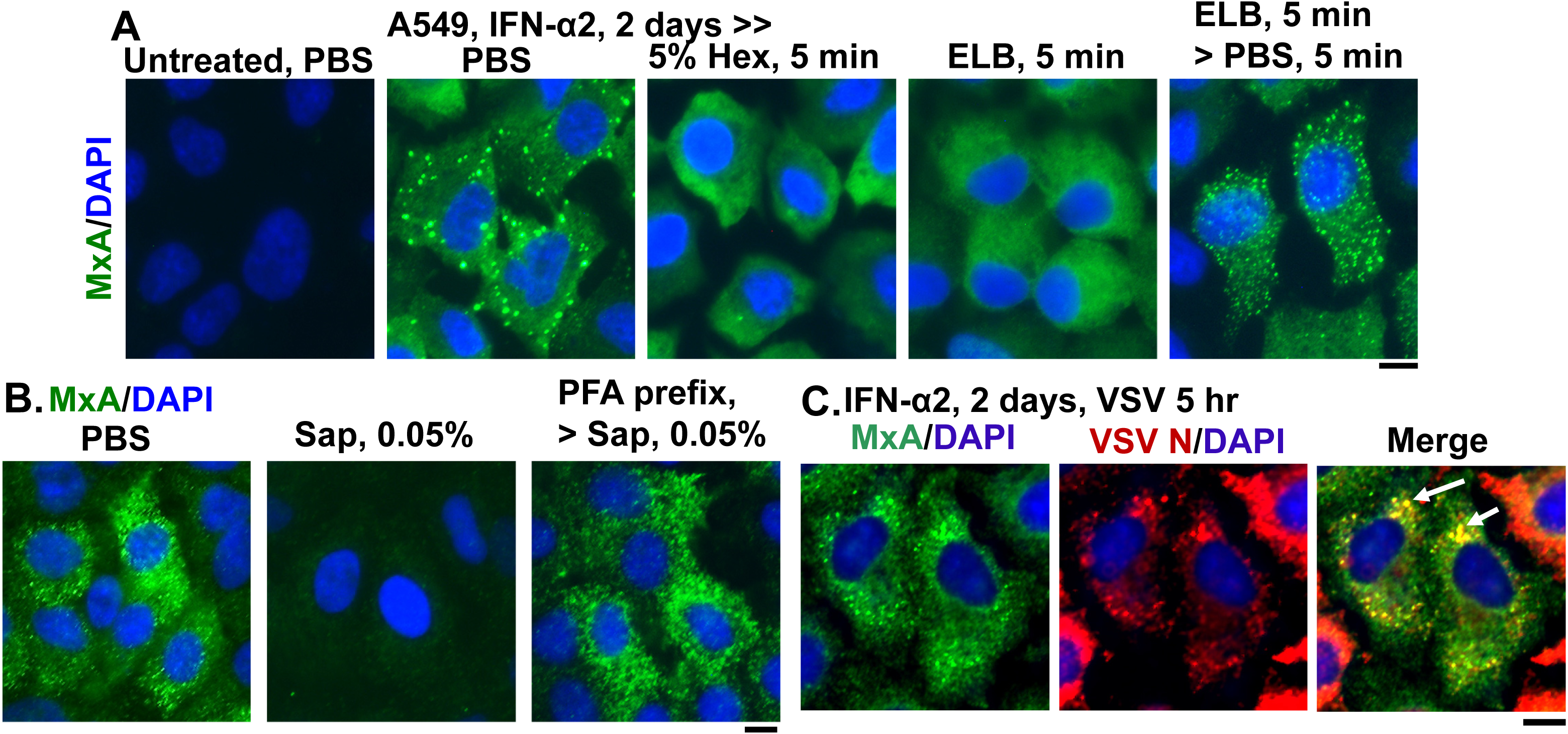
Endogenous MxA granules in IFN-*α*2-treated A549 cells have properties of phase-separated biomolecular condensates. Panel A, A549 cells in culture (in 35 mm plates) without or with exposure to IFN-α2 (20 ng/ml) for 2 days were washed with PBS and then fixed (using in 4% PFA), or exposed to 5% 1,6-hexanediol in PBS for 5 min or to ELB for 5 min or ELB for 5 min and then PBS for 5 min prior to fixation (using 4% PFA). All cultures were then permeabilized (saponin-digitonin buffer) and immunostained for MxA. Panel B, IFN-α2-treated cultures (20 ng/ml, 2 days) were fixed (4% PFA, 1 hr) after a PBS wash, or were first exposed for 10 min to PBS containing 0.05% saponin, and then fixed (2% PFA for 10 min at room temperature) or first fixed with PFA (2% 10 min, room temperature), followed by saponin exposure (0.05% in PBS for 10 min). All cultures were then immunostained for MxA. Panel C, IFN-α2-treated cultures (10 ng/ml, 2 days) were infected with VSV (moi >10 pfu/cell) (44, 46) for 5 hr, fixed and immunostained for MxA (green) and for VSV N (red). Arrows: cells showing condensates with both MxA and VSV N protein. All scale bars = 10 μm.

**Figure 3.**
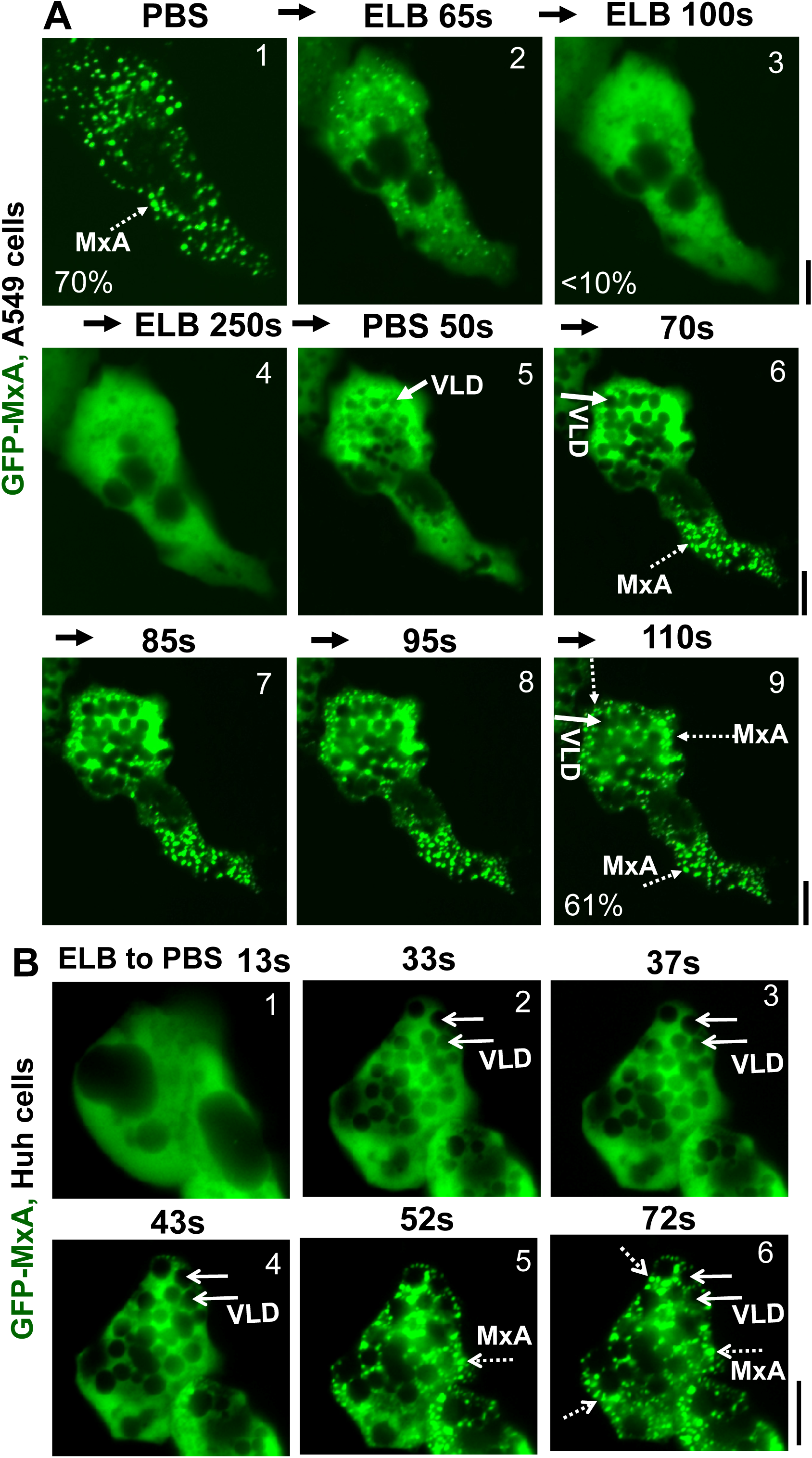
Marked accumulation of vacuole-like dilations (VLDs) in cells preceding isotonicity-driven reassembly of GFP-MxA condensates. Panel A, Live-cell time-lapse images (time designated in seconds) of an A549 cell showing rapid disassembly of GFP-MxA condensates upon exposure to hypotonic ELB, and rapid reassembly of GFP-MxA into new condensates upon shift up to isotonic PBS (broken white arrows). Solid white arrows point to VLDs which accumulate in the cytoplasm prior to GFP-MxA condensate reassembly. The % numerals in respective panels are an estimate of the fraction of total cellular GFP-MxA in condensates. Panel B, Live-cell time-lapse images (time designated in seconds) of a GFP-MxA expressing Huh7 cell previously shifted to hypotonic ELB (thus the GFP-MxA is dispersed) showing rapid reassembly of GFP-MxA into new condensates upon shift up to isotonic PBS (broken white arrows). Solid white arrows point to VLDs which accumulate in the cytoplasm prior to GFP-MxA condensate reassembly. Scale bars = 20 µm.

Fig. 2 summarizes data showing that cytoplasmic structures derived from endogenous MxA in IFN-treated A549 lung cells, had key properties previously observed for GFP-MxA biomolecular condensates (44). The endogenous MxA condensates were disassembled rapidly by exposure of cells to 1,6-hexanediol (5%), to hypotonic ELB and also reassembled upon subsequent exposure to isotonic PBS (Fig. 2A). The endogenous MxA condensates required an intact plasma membrane for their integrity in that they were disassembled by exposure of cells to PBS containing saponin (0.05%)(Fig. 2B). This observation that cell integrity was required for maintenance of MxA higher-order structures in the cytoplasm places limitations on interpretation of prior data on MxA oligomerization derived from previous solution-based analyses (40-42, 49-53). Indeed, as previously shown for stress granules and processing bodies (P-bodies)(54–56), prefixing of cells with 2% PFA for 10 min blocked the subsequent dissolution by saponin of MxA cytoplasmic structures (Fig. 2B). Moreover, Fig. 2C shows that the VSV N protein associated with condensates of endogenous MxA in A549 cells. Overall, the data in Figs. 1 and 2, taken together confirm that many of the properties observed previously for exogenously expressed GFP-MxA in Huh7 hepatoma cells, were also observed for cytoplasmic structures formed by endogenous MxA in IFN-α-treated A549 lung cells.

### Accumulation of plasma membrane-derived VLDs precedes isotonicity-induced GFP-MxA condensate reassembly

Due to practical advantages of fluorescence microscopy studies in live cells, investigations of MxA condensate disassembly and reassembly have used GFP-tagged human MxA constructs transiently expressed in A549, A549-hACE2, Huh7 and Hep3B cells at expression levels which match that of endogenous MxA in IFN-treated cells (see Fig. 2A in ref. 44 for one comparison). GFP-MxA expression in such cells led to the appearance of 70-90% of the MxA in condensates (Figs. 3A, and Fig. 8A, B). Fig. 3A illustrates time-lapse images of the same A549 live cell showing the rapid disassembly (to <10% of GFP-MxA in condensates) and then reassembly of GFP-MxA condensates (up to 70% of GFP-MxA in condensates) in response to hypotonicity (ELB) and isotonicity (PBS) respectively. Fig. 3B illustrates an example of GFP-MxA condensate reassembly in a live Huh7 cell following shift from ELB (hypotonic) to PBS (isotonic). Two features in these data stand out: (i) that the reassembled condensates were different from the original condensates, and (ii) that isotonicity-induced rapid reassembly of condensates was preceded by an even more rapid appearance of fluorescence-dark spaces vacuole-like in the cytosol. The latter vacuole-like dilated (VLD) spaces have been reported beginning over 2-3 decades ago as part of the recovery of cells from hypotonic stress and comprise the internalization of plasma membrane large vesicles (7–13). It is noteworthy from the data in Fig. 3 that the new reassembled GFP-MxA condensates were confined to the reduced cytosolic space located between VLDs. The data are consistent with the possibility that GFP-MxA disassembly under hypotonic conditions and subsequent reassembly under isotonic conditions may represent a response to cytosol “uncrowding” and subsequent “crowding.”

Supporting evidence for the above organellar interpretation was obtained by the observations that in cells doubly expressing GFP-MxA condensates and soluble red fluorescent protein (RFP) hypotonic disassembly was accompanied by a mixing of the green and red fluorescence, and subsequent isotonicity-driven GFP-MxA condensate reassembly was confined to RFP-containing inter-VLD spaces (Fig. 4). As expected, the VLD spaces were devoid of cytosolic RFP (Fig. 4), consistent with the formation of VLDs by engulfment of extracellular medium during internalization of the plasma membrane (7–13). Evidence that such VLDs were indeed lined by the plasma membrane was obtained using Alexafluor A594-tagged cholera toxin B (CTB) which binds the plasma membrane (57) in ELB-treated cells, and is then rapidly internalized into VLDs when cells were shifted to isotonic medium (Fig. 5A and 5B). In cells in hypotonic ELB, the disassembled GFP-MxA was distributed in the cell contained within boundaries demarcated by CTB-red-marked plasma membrane (Fig. 5B, top row). Subsequent GFP-MxA condensate reassembly took place outside of the CTB-red lined VLD spaces (Fig. 5B, bottom row).

**Figure 4.**
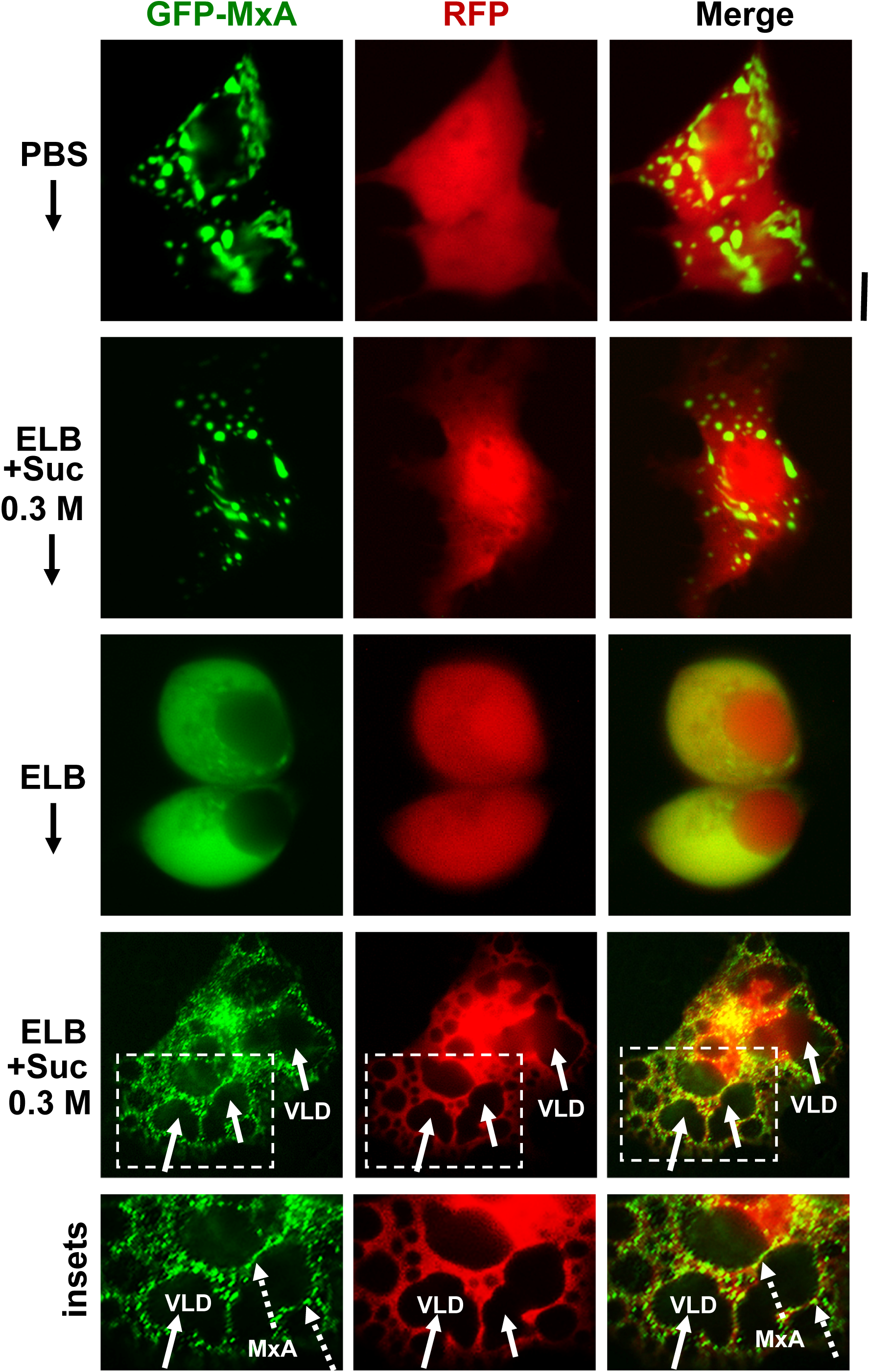
Isotonicity-driven reassembly of GFP-MxA condensates occurs in cytosolic spaces in-between crowded VLDs. Huh7 cultures were co-transfected with vectors for GFP-MxA and soluble RFP, and cells expressing both fluorescent proteins were imaged under respective changes of the culture medium as indicated (5-10 min per change). Boxed insets are shown at higher magnification in the lower-most row. Solid arrows point to VLDs, broken arrows to freshly reassembled GFP-MxA condensates. Scale bar = 10 µm.

**Figure 5.**
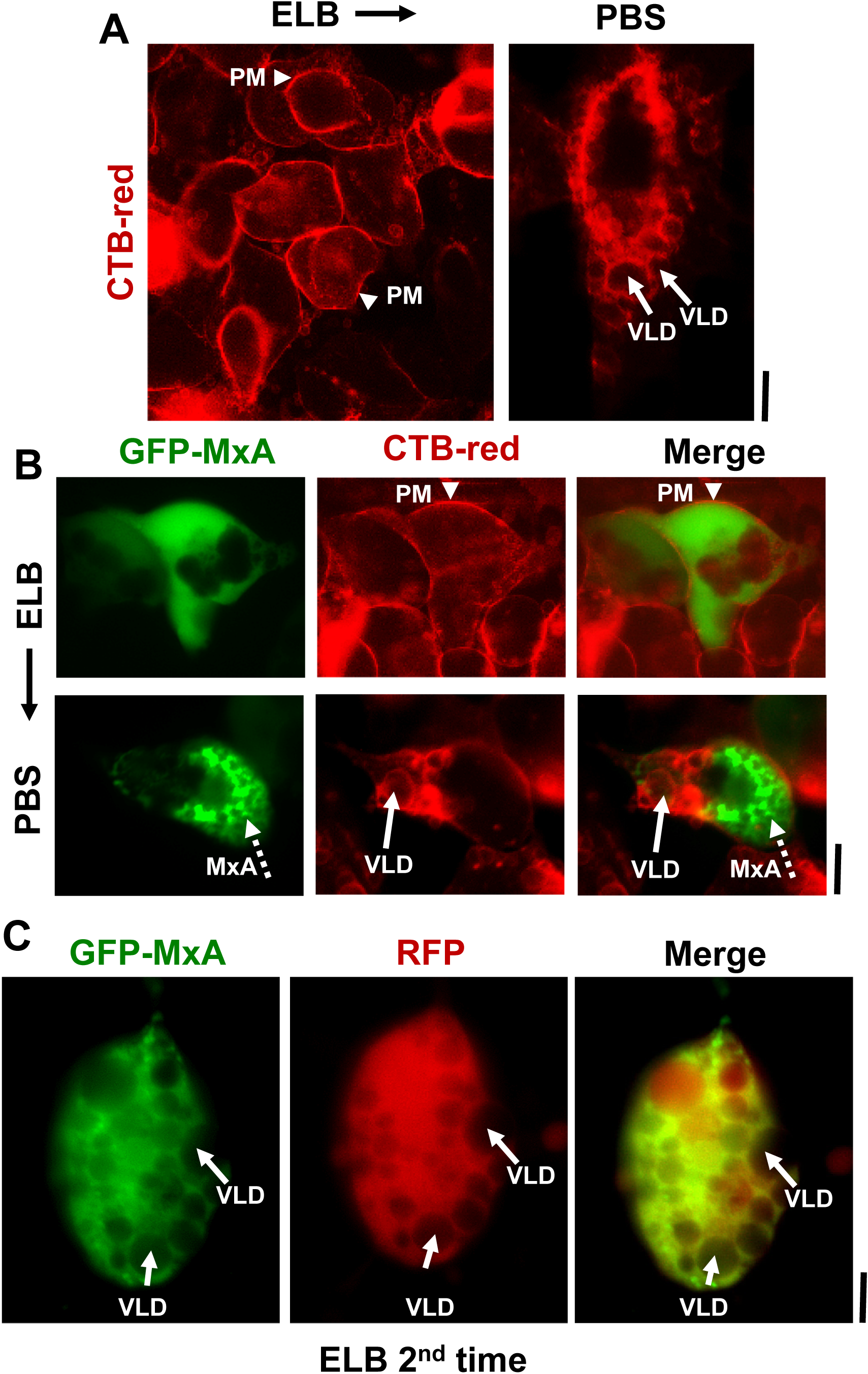
Isotonicity-driven VLDs represent internalized plasma membrane but survive a second hypotonic challenge. Panel A, Live Huh7 cells swollen in hypotonic ELB were labelled with Alexafluor 594-tagged cholera toxin-red (CTB-red; 10 µM)(left panel), imaged, and were then shifted to isotonic PBS (5 min)(right panel) and reimaged. Arrow heads point to plasma membrane (PM), solid white arrows to VLDs. Panel B, Live Huh7 cells expressing GFP-MxA (green) first swollen in hypotonic ELB, then labelled with CTB-red for 5 min (top row), were shifted to isotonic PBS and reimaged 2-3 min later (bottom row). Arrow head, plasma membrane (PM). Solid white arrows point to VLDs, broken white arrows to reassembled GFP-MxA condensates. Panel C. Live Huh7 cell expressing both soluble RFP and newly reassembled GFP-MxA condensates (as at the end of the experiment in Fig. 4; lower-most panels), were re-exposed to hypotonic ELB for a second time. Such cells, imaged 2-3 min later as in this panel, showed newly disassembled GFP-MxA despite the continued presence of a cytoplasm crowded with VLDs (solid white arrows). All scale bars = 10 µm.

We have previously reported that ELB-driven disassembly and isotonicity-driven reassembly of GFP-MxA condensates could be repeated up to 3 times (Figs. 9A and 10A in ref. 44). Curiously, the previous data showed that cells undergoing a second or third cycle of ELB-driven GFP-MxA disassembly continued to exhibit VLD-like dark spaces in the cytoplasm despite the disassembly and dispersal of GFP-MxA (Figs. 9A and 10A in ref. 44). We have now confirmed the earlier findings by the observation that a second cycle of ELB-driven GFP-MxA disassembly can indeed take place in cells that continue to exhibit marked accumulation of VLDs in the cytoplasm (Fig. 5C). Thus, while cytosolic “crowding” subsequent to extensive VLD internalization needs to be considered as a contributor to mechanisms driving MxA condensate reassembly, VLD formation *per se* cannot be the sole mechanism regulating this process.

GFP-MxA stability in cells under isotonic conditions, condensate disassembly in cells exposed to hypotonic ELB and subsequent reassembly cells shifted to isotonic medium was not affected by inclusion of ZnSO_4_ (200 µM)(Zn has been reported by some investigators to stabilize condensates; 58), or CaCl_2_ (200 µM) or the calcium chelator EGTA (1 mM) in the respective buffers (data not shown). Moreover, isotonicity-driven GFP-MxA condensate reassembly was selective to MxA in that condensates of GFP-tagged SARS-CoV-2 virus nucleocapsid (N) protein did not exhibit this reassembly in isotonic PBS (Suppl. Fig. 1). As has been reported previously by others in other cell types (59–62), we observed that approximately 30-40% of SARS-CoV-2 N-GFP expressed in transiently transfected A549 cells was located in condensates as judged by their disassembly in 1,6-hexanediol (Suppl Fig. 1A). These condensates also disassembled when cells were exposed to hypotonic ELB (Suppl. Fig. 1B; to <10% in condensates). However, CoV-2 GFP-N did not reassemble back into condensates in cells shifted to isotonic PBS (Suppl. Fig. 1B). That there was a clear distinction between condensates formed by MxA and those formed by CoV-2 N is also evident in the data in Suppl Fig. 2, which shows that these two proteins formed completely distinct condensates in the cytoplasm of cells transiently co-transfected with both expression vectors (tagged either with GFP or HA in different combinations). Nevertheless, parenthetically, transient expression of GFP-MxA *per se* in Vero cells exhibited an antiviral activity towards SARS-CoV-2-mCherry virus (P. B. Sehgal, unpublished data).

### Spontaneous GFP-MxA condensate reassembly under moderate hypotonic stress

ELB was used in the preceding experiments as a hypotonic buffer because it has been used over the decades by investigators to swell cells prior to mechanical breakage (such as Dounce fractionation protocols; see 47 for one example); ELB (∼40-50 mOsm) is significantly hypotonic compared to isotonicity (∼330 mOsm). Cells kept in ELB for an hour or more showed persistent disassembly of GFP-MxA structures and eventually lifted off the culture dish (not shown).

The data in Fig. 6 summarize the ability of more moderate levels of hypotonicity to disassemble GFP-MxA condensates. In this experiment, cultures were exposed to hypotonic medium diluted with distilled water to give 1/4, 1/3, 1/2 and 3/4 strength tonicity (range 90-175 mOsm). The data in Fig. 6A show moderate hypotonicity-dependent disassembly of GFP-MxA condensates over the next 9-10 min. Quantitation of these data was carried out using the Fourier transformation and size-filter approach (in Image J)(as illustrated in Fig. 7B), The numerical estimates of the extent of disassembly (Fig. 6B) show detectable disassembly in 3/4 tonicity medium, with clearcut disassembly apparent in half-tonicity medium by 2-3 min. Significant disassembly to 10-20% residual GFP-MxA in condensates was observed in cells exposed to one-third or one-fourth tonicity medium in less than 10 min (Fig. 6B).

**Figure 6.**
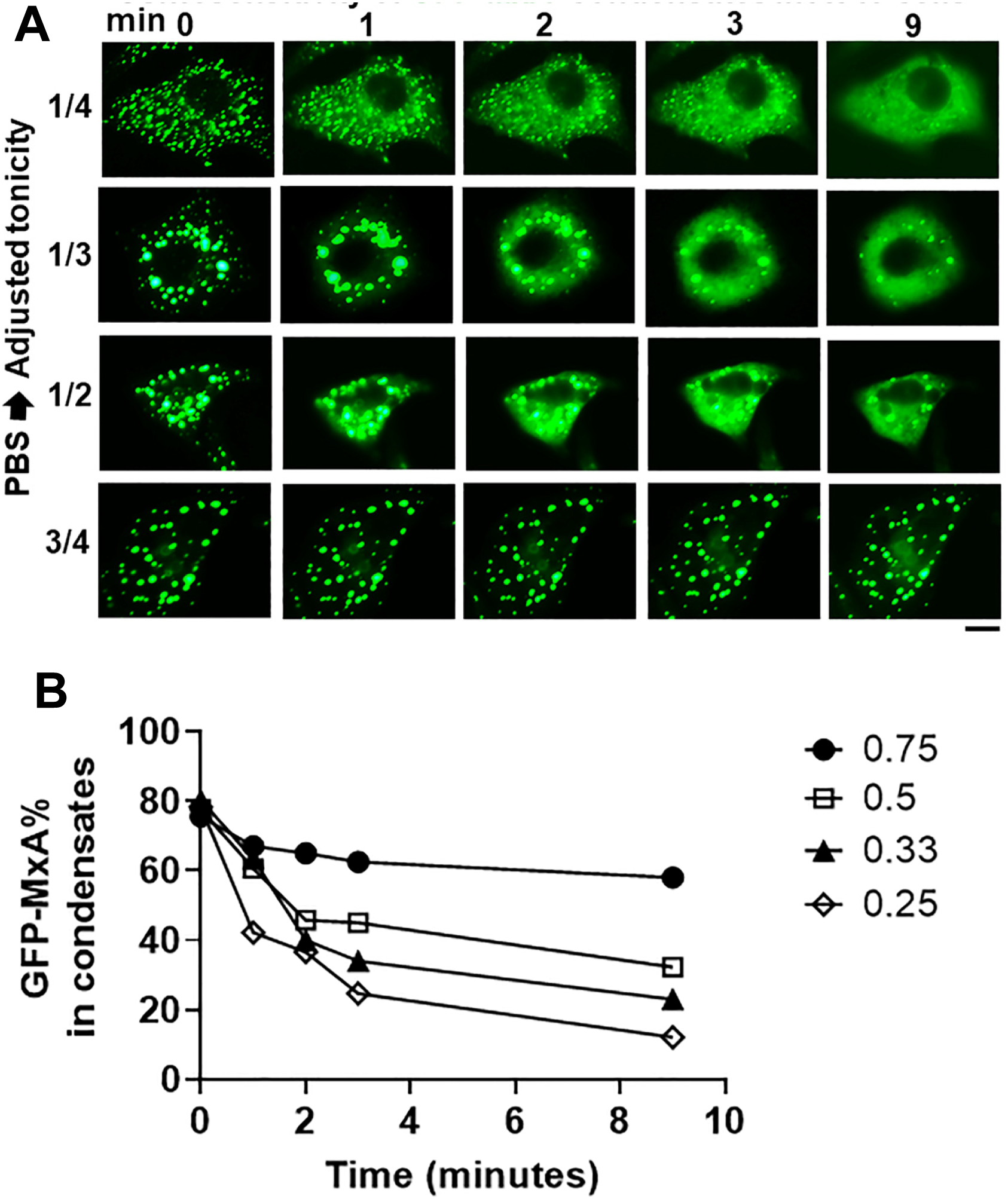
Hypotonicity-concentration dependence of the disassembly of GFP-MxA condensates. Oanel A, Cultures of live A549 cells expressing GFP-MxA condensates were first imaged in isotonic PBS, and then shifted to PBS diluted to 1/4, 1/3, 1/2, and 3/4 strength isotonicity using water. Individual cells in respective cultures were imaged in a time-lapse mode. Scale bar = 10 µm. Panel B, Quantitation of the extent of GFP-MxA (as % of total in the cell) in condensates in the images shown in Panel A. This quantitation was carried out using a Fourier filter in Image J to subtract objects of small radius (as illustrated in Fig. 7B).

In as much as phase contrast microscopy showed that confluent A549 cultures remained intact for up to 5 hr in hypotonic medium in the range 1/4, 1/3, 1/2 isotonicity, we extended the time of observation of GFP-MxA structures in cells exposed to moderate hypotonic medium for up to 60 min. Much to our surprise, in such cells, the disassembled GFP-MxA *spontaneously* reassembled into new condensates by 7-15 min (Fig. 7A and B). This spontaneous reassembly was preceded by the appearance of VLDs in such cells (Fig. 7A). Quantitation of the extent of GFP-MxA in condensates vs diffuse state in a cell was carried out using the Fourier-based “minimum” filter in Image J to subtract small objects such as condensates from images (Fig. 7B). Supplemental Movie 1 and Supplemental Movie 2 show time-lapse movies (5 sec/frame) of the disassembly of GFP-MxA condensates in an A549-hACE2 and Huh7 cell respectively shifted to one-third isotonic medium, and the subsequent spontaneous reassembly of GFP-MxA into new condensates in such cells 10-15 min later. These live-cell movies in the two cell types again show the appearance to the dark GFP-free VLD spaces prior to the reassembly of GFP-MxA into condensates. Remarkably, the spontaneous reassembly of MxA condensates under moderate hypotonicity conditions was also confirmed for endogenous MxA condensates/granules (Fig. 7C).

**Figure 7.**
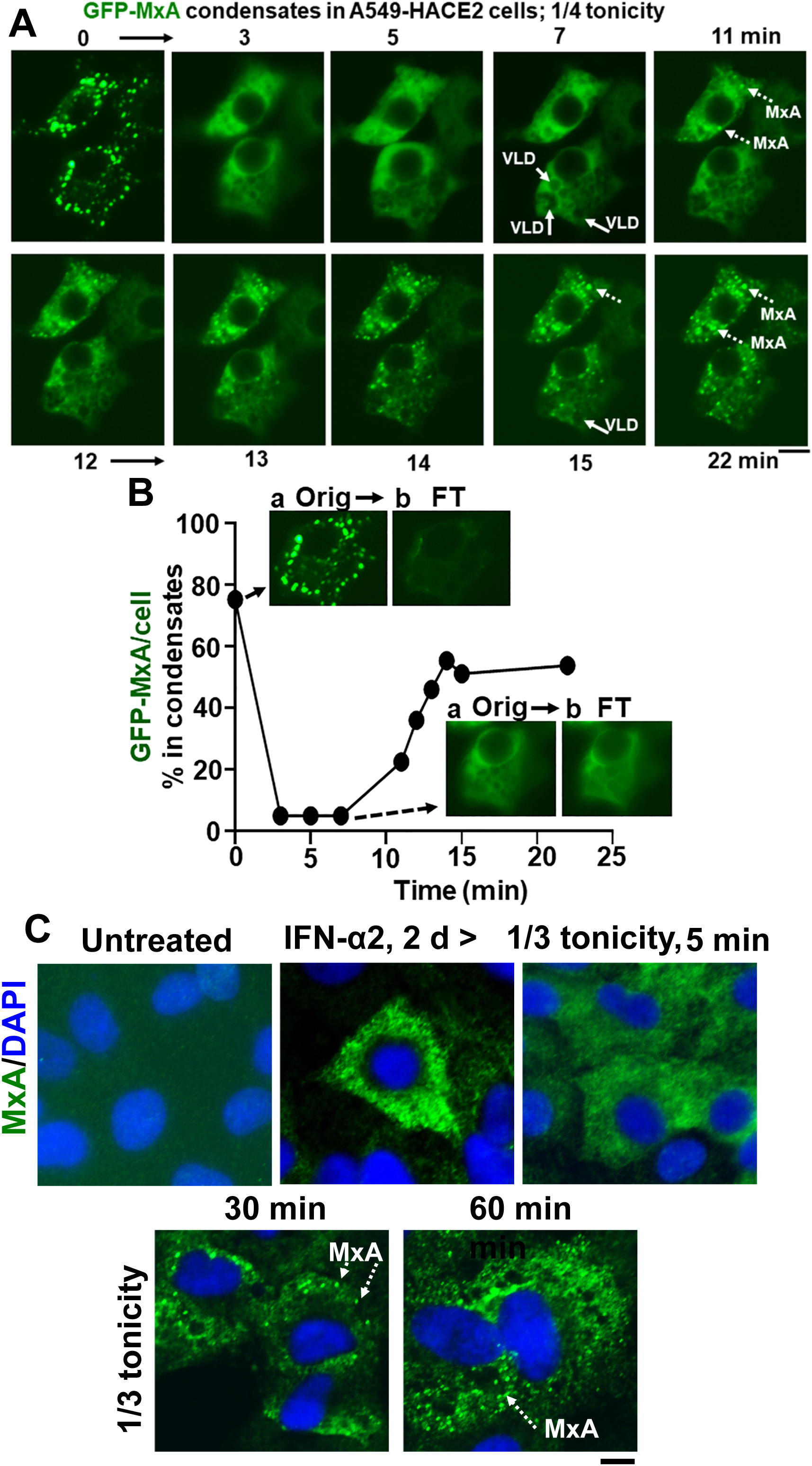
Spontaneous reassembly of MxA condensates despite continued moderate hypotonic stress. Panel A, Cultures of live A549 cells containing GFP-MxA condensates were exposed to 1/4 strength hypotonic stress (1 part full culture medium diluted with 3 parts distilled water) an the cells imaged in a time-lapse mode for 25-20 minutes. The panel illustrates two representative cells displaying rapid disassembly of GFP-MxA condensates and then spontaneous reassembly beginning 11 min later. Solid arrows point to VLDs, broken arrows to spontaneously reassembled GFP-MxA condensates. Panel B, shows quantitation of GFP-MxA in condensates (as % of total) in the lower cell in images in Panel A. Insets show the process used to subtract condensates from respective images using a Fourier filter in Image J (insets a show the original image and insets b the filtered image with the condensates subtracted; these can be further quantitated using Image J to derive total and dispersed GFP values). Orig, original image; FT, filtered image. Panel C, Replicate IFN-α2 (20 ng/ml, 2 days) treated cultures were exposed to 1/3 tonicity culture medium for 5, 30 and 60 min, then fixed using 4% PFA (in 1/3 tonicity PBS), followed by immunostaing for endogenous MxA. Broken white arrows point to newly spontaneously reassembled condensates of endogenous MxA. All scale bars = 10 µm.

The discovery of *spontaneous* reassembly of GFP-MxA condensates in cells under moderate hypotonic conditions, allowed for testing the contribution of various biochemical mechanisms towqrds GFP-MxA condensate formation. Fig. 8 summarizes an experiment in which dynasore (100 µM) was observed to reduce the extent of reassembly – an observation consistent with the possibility that plasma membrane internalization (such as to form VLDs) could drive MxA reassembly. In contrast, dynasore (100 µM) had little effect on the massive VLD development observed in switching cells from the markedly hypotonic ELB medium to isotonic PBS, and the subsequent very rapid (within 30-60 sec) GFP-MxA reassembly as in Fig. 3 (data not shown).

**Figure 8.**
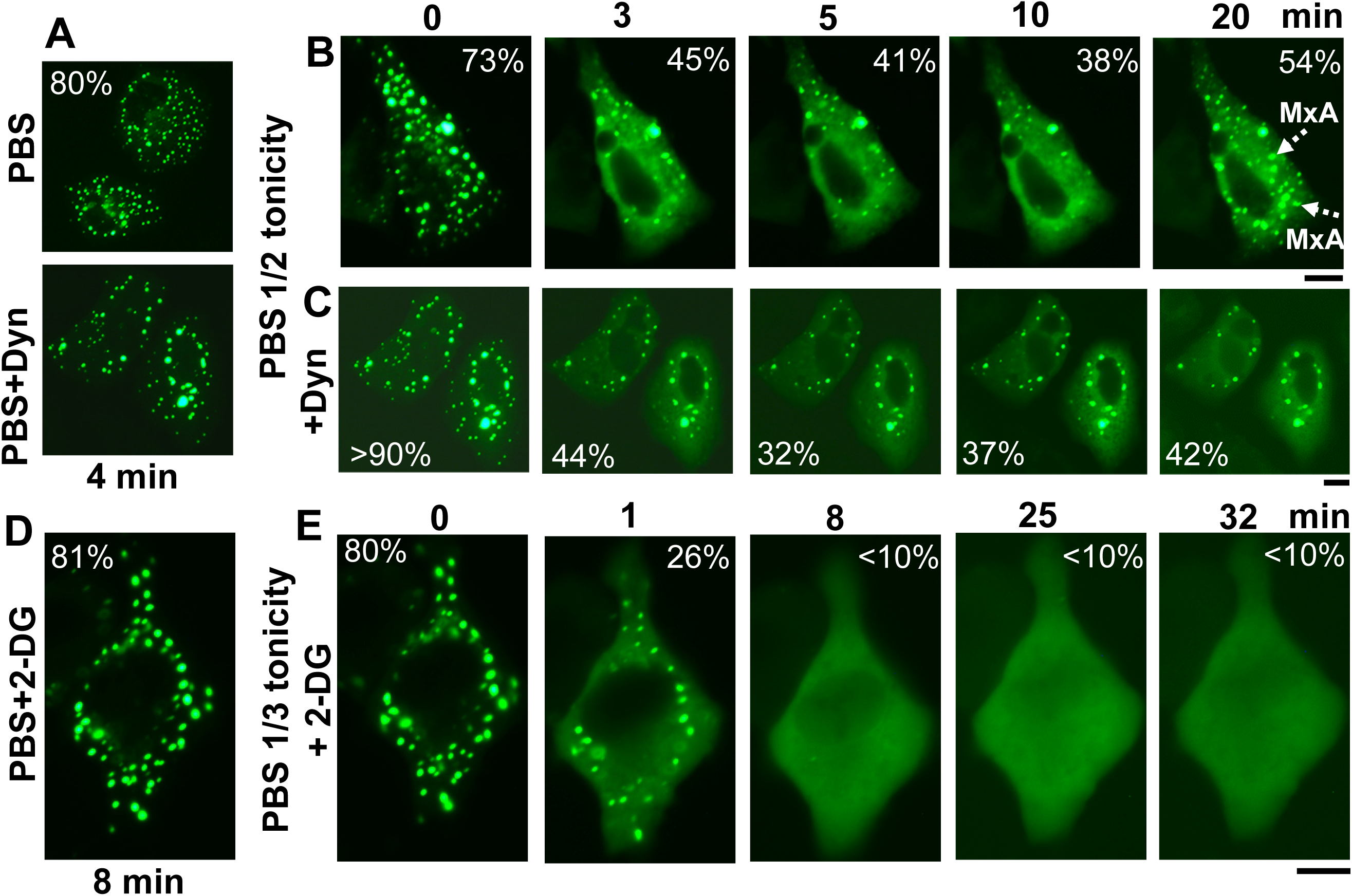
Spontaneous reassembly of GFP-MxA condensates is inhibited by dynasore and 2-deoxyglucose (2-DG). Panels A, B and C, Cultures of A549 cells expressing GFP-MxA were shifted to PBS and then exposed dynasore (Dyn; 100 µM in PBS) for 4 min, followed by a shift to one-half strength PBS without or with inclusion of dynasore (100 µm). Panel illustrates a compilation of respective time-lapse images; numerals in each panel indicate % of GFP-MxA in condensates (evaluated as per the Fourier filter method depicted in Fig. 7B). Broken white arrows point to newly spontaneously reassembled GFP-MxA condensates. Panels D and E, Cultures were treated with PBS supplemented with 2-DG (10 mM) for 8 min, and then shifted to one-third strength PBS with inclusion of 2-DG (10 mM).

The slower spontaneous reassembly assay (as in Fig. 7) was used to investigate the effects of various inhibitors previously shown to stabilize condensates or affect water or ion trafficking. The inclusion of 2-deoxygluxcose [2-DG; 10 or 20 mM; this inhibits ATP generation (63)] in full-strength PBS did not affect GFP-MxA condensate structures (Fig. 8D). The inclusion of 2-DG in 1/3rd strength PBS did not affect GFP-MxA disassembly (Fig. 8E), However 2-DG inhibited condensate reassembly in 1/3rd tonicity medium (Fig. 8E), suggesting that this spontaneous reassembly was energy dependent.

The dual kinase phosphorylation inhibitor GSK-626616 (1 µM, which inhibits condensate disassembly during mitosis; 64), the condensate “hardening” agent cyclopamine hydrate [which is an antiviral against respiratory syncytial virus (RSV) shown to “stabilize” RSV condensates against hypotonic disassembly (in response to 1/10 strength culture medium; 65) when used at 10 µM], thapsigargin (a calcium-ion channel inhibitor; 20 µM), inhibitors of various aquaporin channels (66) [tetraethylammonium (TEA, up to 10 mM), *N*-1,3,4-thiadiazol-2-yl-3-pyridinecarboxamide (TGN-020; 10 µM), copper sulfate (100 µM), singly or in combination, had little effect on stability of GFP-MxA condensates in cells kept in isotonic medium, their disassembly in moderately hypotonic medium (1/4th strength tonicity) or subsequent spontaneous reassembly (data not shown).

### Antiviral activity of GFP-MxA against VSV survives osmotic disassembly/reassembly

We monitored the replication of VSV at the single-cell level (by immunofluorescence for VSV N) in cultures transiently transfected with a GFP-MxA construct and then with VSV at moi >10 pfu/cell followed by fixation 4 or 5 hr after infection which is near the peak of VSV N expression in infected cells (44, 67, 68). Because we wished to evaluate the effect of disassembly and reassembly of GFP-MxA condensates lasting several minutes to a few hours on VSV replication soon after initiation of the infection, the approach used required a single-cycle replication assay under conditions in which >90% of cells in a culture were infected (moi > 10 pfu/cell), and not a low-multiplicity multiple-cycle virus assay. Moreover, VSV N protein expression was evaluated by immunofluorescence (in red) at the single-cell level in GFP-MxA expressing and GFP-MxA-free cells located side-by-side in the same culture. Thus, all cells (without or with GFP-MxA expression) were exposed to the same virus and the same osmotic manipulation, and thus the cells without GFP-MxA expression (by subsequent fluorescence microscopy in green) served as a control for the effect (if any) of osmotic manipulation on VSV replication *per se* (quantitated by red immunofluoresence).

Data in Suppl Fig. 3, left most culture, validate this approach - GFP-MxA expressing cells and showed a marked reduction in VSV N immunofluorescence (also see refs. 44 and 46 for additional examples). The possibility that cellular edema (hypotonicity-induced GFP-MxA disassembly) might affect antiviral activity was investigated in two ways. In one approach, cells the infected GFP-MxA expressing cultures were subjected to a 5 min cycle of ELB and then returned to full culture medium at 1, 2 or 3 hr after infection, and the antiviral activity assessed at the single-cell level at 4 hr after infection (Suppl Fig. 3). In the second approach, cultures were switched to one-third tonicity medium 45 min after the start of infection, fixed at 5 hr after start of infection, and expression of VSV N evaluated in GFP-MxA-negative and in GFP-MxA positive cells. In both instances, the disassembly and reassembly of GFP-MxA (as expected) was verified by live-cell imaging of the same cultures (not shown).

The data in Suppl Fig. 3, middle and right-most cultures, show that one 5-min cycle of disassembly/reassembly of GFP-MxA at 1 hr or 3 hr after the beginning of VSV infection had little effect on the antiviral activity. In a similar experiment, one 5-min cycle of disassembly/reassembly at 2 hr after the start of infection also had little effect on antiviral activity (not shown). Fig. 9A and 9B show that continued maintenance of cells in one-third isotonic medium also had little effect on the antiviral activity of GFP-MxA. Remarkably, many cells showed the co-localization of VSV-N with the reassembled GFP-MxA condensates (Fig. 9A). Overall, the antiviral activity of GFP-MxA against VSV largely survived this edema-like hypoosmolar stress.

**Figure 9.**
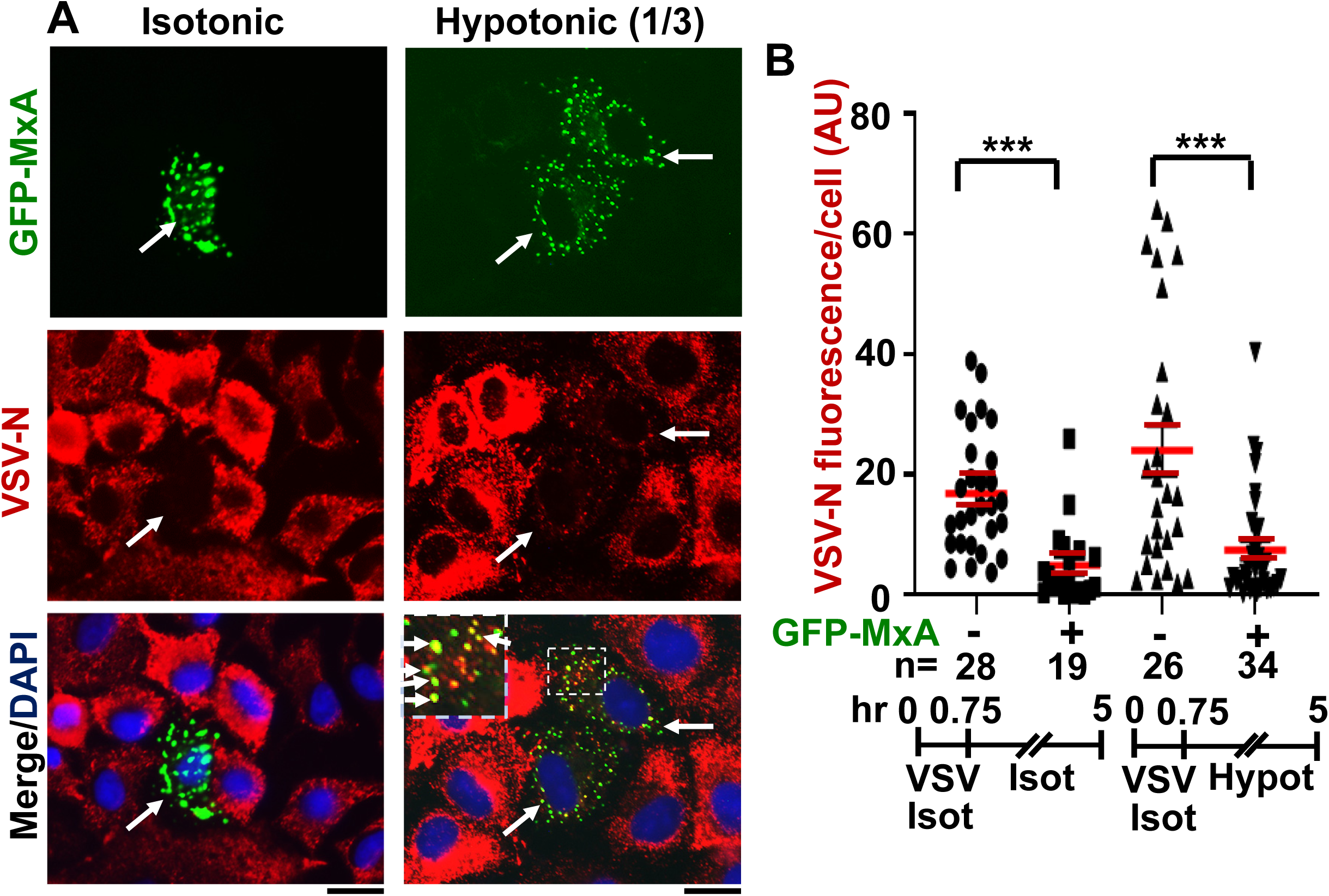
Antiviral activity of GFP-MxA against VSV survives hypotonic stress. **Panel A,** A549 cells were transfected with pGFP-MxA and 2 days later were exposed to VSV (moi >10 pfu/cell) for 45 min in isotonic serum-free DMEM (Isot). Cultures were then washed and maintained for the next 4.25 hr in regular medium (Isot) or one-third strength medium (Hypot). The cultures were then fixed and imaged for GFP-MxA (green) and for VSV-N (immunostained in red). Solid white arrows: GFP-MxA expressing cells with reduced VSV-N. Inset: GFP-MxA condensates which incorporate VSV N (white arrows). Scale Bar = 20 μm. Panel B, quantitation of VSV N per cell in GFP-MxA positive (+) or negative (-) individual cells in the same culture kept without or with one-third hypotonic stress in the experiment in Panel A. n = number of cells evaluated per group in this experiment; horizontal red lines within each group indicate Mean ±SE. Statistical significance was evaluated using one-way ANOVA (Kruskal-Wallis with Dunn’s post-hoc test for multiple comparisons); *** *P* <0.001.

## Discussion

The present study highlights a dramatic property of phase-separated biomolecular condensates of the human antiviral MxA protein – rapid reversible sensitivity to cellular responses to hypoosmolar changes in the extracellular fluid (Fig. 10). Lung- and liver-derived cells exposed to even mild hypotonicity exhibited rapid disassembly of MxA condensates within 1-3 min. Remarkably, the dispersed MxA reassembled into new condensates either upon subsequent shifting cells to isotonic medium (induced reassembly) or even spontaneously in cells maintained in moderately hypoosmolar conditions (Fig. 10B). Isotonicity-driven or even spontaneous MxA condensate reassembly was preceded by the accumulation of plasma membrane-lined VLDs in the cytoplasm (Fig. 10). The biochemical basis for a relationship between VLD dynamics and MxA condensate formation, such as “crowding” of the liquid phase of the cytosol by VLDs, remains intriguing. Importantly, the antiviral activity of MxA against VSV, a virus which replicates exclusively in the cytoplasm, survived osmotic disassembly/reassembly. As a technical issue, the observation that cell integrity was required for maintenance of MxA higher-order structures in the cytoplasm places limitations on interpretation of prior data on MxA oligomerization derived from numerous previous solution-based analyses.

**Figure 10.**
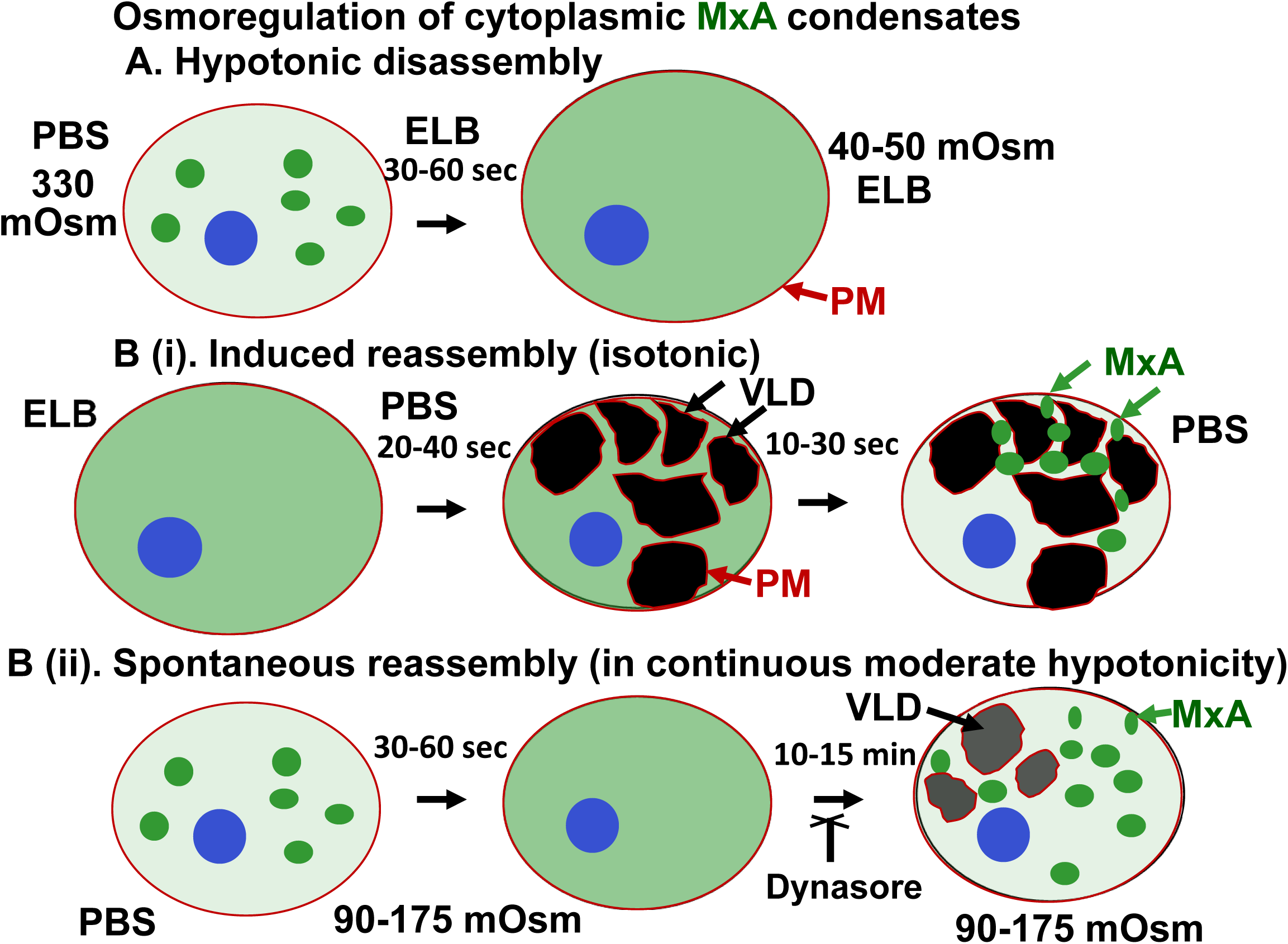
Schematic summary of osmoregulation of cytoplasmic MxA condensates. ELB, hypotonic erythrocyte lysis buffer; PBS, phosphate-buffered saline; PM, plasma membrane (displayed using CTB-red as in Fig. 5); VLD, vacuole-like dilatation.

We previously reported that endogenous MxA induced by exposure of primary human pulmonary arterial endothelial cells in culture to IFN-α was present in the cytoplasm in granular structures (43, 69). In the present study we provide evidence showing that cytoplasmic structures produced by endogenous MxA induced by IFN-α in lung derived cells had properties similar to biomolecular condensates formed by exogenously expressed GFP-MxA – disassembly by 1,6-hexanediol, disassembly in cells exposed to hypotonic ELB, and reassembly upon isotonic shift; and incorporation of the viral (VSV) nucleoprotein in MxA condensates (Fig. 2). An intact plasma membrane was necessary to maintain MxA condensate integrity. Puncturing the plasma membrane with saponin at a low concentration (0.03-0.05%) led to disassembly of almost all the condensates produced by endogenous MxA (Fig. 2B), and most of those produced by exogenously expressed GFP-MxA (44). With both endogenous MxA (Fig. 2B) and GFP-MxA (44), saponin caused a net loss of MxA from the cytoplasmic compartment (into the culture medium). However, as with data previously obtained by other investigators in studies of stress granules and P-bodies (54–56), pre-fixing of cells with paraformaldehyde (2% at room temperature for 10 min) stabilized MxA condensates to a detergent exposure (Fig. 2B). Thus, prior attempts to identify cellular proteins which associated with MxA using protocols involving cell breakage likely provide incomplete answers (41–42).

MxA is a dimeric protein which readily multimerizes into larger linear and ring-shaped assemblies in solution (41-42, 49-53). The globular GTPase is hinged to a stalk region consisting of 4 α-helicies linked one to the other by disordered loops (L1 to L4). The L4 loop contains a stretch of 4 Lys residues important for multimerization and antiviral activity. It is already clear that MxA can readily form protein clusters (higher-order oligomers) in solution (49–53). Although the L4 loop represents an intrinsically disordered region, the structural bases for the formation of phase-separated MxA condensates are incompletely understood. Prior cell-free solution-based data of the oligomerization of MxA are the reverse of what we now report for MxA in intact cells. Specifically, Haller and colleagues reported that in cell-free solutions recombinant MxA formed large linear and ring-shaped oligomers in low salt conditions (50 mM NaCl); these persisted at 150 mM NaCl but were disrupted into small structures at 300 mM NaCl (49–52). Oligomerization was further enhanced in the presence of GDP-Al fluoride or GMP-PCP (49–52). The previously published studies in cell-free solutions emphasizing MxA oligomerization under low-salt conditions (50 mM NaCl) are consistent with electrostatic interactions driving oligomerization in solutions of recombinant MxA. However, in intact cell studies, we now report that hypotonic exposure of cells disrupts MxA condensates, while isotonic (and even hypertonic) exposure of cells reassembles condensates. Recent biophysical insights (56, 70, 71) which suggest a continuum of “fuzzy interactions” (low-affinity with high/variable stoichiometry) depending on cytosol context between protein monomers with a multivalent tendency, with or without the participation of other protein components or nucleic acid polymers, to generate “protein clusters” extending up in size to the formation of higher-ordered structures and even phase-separated condensates, may apply to human MxA in the cell cytoplasm. Our observations that puncturing the cellular plasma membrane using low-concentration of saponin (0.03 to 0.05%) leads to a major loss of condensates formed by GFP-MxA (44) and endogenous MxA (Fig. 2B) suggests that the behavior of recombinant MxA in solution (as monomers, dimers and tetramers and even assembly of higher-order oligomers in low-salt solutions) as elucidated over the last two decades (41-42, 49-52) may not be fully applicable to the physiological behavior of MxA protein clusters/condensates in the protein-crowded cytosol of the intact cell (disassembly in low-salt media). This limitation of studies of protein oligomerization in free solution is likely applicable to other proteins – our data suggest that the transcription factor STAT3 (92 kDa) also forms variably sized clusters (of size in the range from 200 KDa to 2 MDa) in the cytosol and can also form phase-separated cytoplasmic and nuclear condensates (47, 48, 72).

The dramatic accumulation of VLDs in the cytoplasm of cells shifted from hypotonic conditions to isotonic conditions has been recognized in numerous studies over the last 3 decades (7–13). In particular, Sheetz and colleagues have investigated the generation of mechanical plasma membrane tension in mast cells and neuronal cells subjected to tonicity changes (8, 11, 12). A decrease in plasma membrane tension (such as when swollen cells are shifted to isotonic medium) leads to marked endocytosis as the cell readjusts its surface membrane area. The endocytosed structures formed included large mesoscale vacuole-like dilations (VLDs) which crowd the cytosolic space. It is noteworthy that in the experiments shown in Fig. 3 and Supplemental Movies 1 and 2, VLD formation preceded the reassembly of GFP-MxA condensates. Thus, cytosolic crowding due to this accumulation of VLDs may represent a mechanism contributing to MxA phase separation and condensate formation. That dynasore, which inhibits clathrin-mediated endocytosis, showed a modest inhibitory effect on MxA condensate reassembly is consistent with a role of plasma membrane internalization in the osmo-mechanical regulation of MxA condensate formation. This mechanism was selective in that there was little reassembly of condensates of the nucleocapsid protein (N-GFP) of SARS CoV-2 virus when cells were cycled from hypotonic conditions to isotonic medium (Suppl. Fig. 1). Thus, different condensates even in the same cell can respond differently to cellular events during osmotic stress. Moreover, the data in Fig. 5C emphasize that accumulation of VLDs in the cytoplasm may not be the only driver for MxA condensate assembly. Fig. 5C (and Fig. 9A and 10A in ref. 44) show that a second round of GFP-MxA hypotonic (ELB) disassembly can occur in cells in the presence of significant VLD accumulation. Thus, the relationship between VLD accumulation and “crowding” in the biophysical sense of removal of water molecules from the liquid phase of the cytosol remains unclear. The possibility that additional biochemical mechanisms participate in the osmoregulation of MxA condensate disassembly/ reassembly remains open. Indeed, MxA has been shown to interact physically with the TRPC-family of stretch receptors (TRPC-1, -3, -4, -5, -6 and -7) and to regulate Ca^++^ signaling (73).

Hyperosmotic stress (HOPS) also promotes reversible condensate formation in human cells by multivalent proteins, especially those associated with P bodies (15–17). The trimeric protein DCP1A, expressed in human cell lines, underwent rapid and reversible condensate formation when cells were shifted from isotonic medium (150 mM NaCl) to hypertonic medium (300 mM NaCl). More generally, proteins with a self-interacting valency ≥2 underwent HOPS (15). The functional consequences included transcription termination defects during osmotic stress because of condensate formation by transcription accessory proteins such as CPFS6 (15). In the case of GFP-MxA, hyperosmotic medium made the condensates more compact and cigar-like in intact cells (44). In contrast, Haller and colleagues have reported that in cell-free solutions 300 mM NaCl caused the truncation of MxA oligomers into smaller structures (49–51).

The relationships between MxA condensate formation and antiviral activity appear to be complex. The structural basis for the antiviral effect of human MxA against both the myxoviruses such as FLUVA (which include a nuclear 5’ cap-snatching step in their replication) and rhabdoviruses such as VSV (which replicate entirely in the cytoplasm) are incompletely understood. MxA is customarily considered to inhibit early transcription of the incoming virus in the cytoplasm (such as when targeting VSV) as well as later steps in virus replication (such as when targeting FLUAV). Cell-free experiments confirm the ability of recombinant MxA to inhibit viral transcriptional activity (40–42, 74). However, it is unclear whether or how monomeric, dimeric or higher-order structures of MxA associate with viral components in mediating this inhibition in intact cells. The present quantitative data show a dynamic equilibrium between MxA amounts in cytoplasmic condensates and in the dispersed mode in IFN-α-stimulated cells or in cells expressing GFP-MxA with 60-80% of the MxA in condensates. These data are consistent with the presence of least some dispersed MxA at all times in most cells. Previous mutational studies of human MxA showed that the GTPase activity was required for most of its antiviral activity (except that against hepatitis B virus) (40-42, 51-53). Inspection of data in the literature also reveals that MxA mutants lacking GTPase activity (and thus antivirally inactive), can still form cytoplasmic condensates (51, 52). Mutations that cause dispersal of MxA in the cytoplasm (e.g. the D250N mutant) lacked antiviral activity (52, 74). However, the R645 point mutant of MxA, which formed larger *cytoplasmic* granules, had the unusual property of inhibiting FLUAV but not VSV, even though the wild-type MxA shows antiviral activity towards both viruses (75). Overall, it appears that the ability to form condensates by MxA may be important but not sufficient for the antiviral activity of Mx proteins.

The subcellular compartment within which Mx proteins are localized also determines their antiviral activity. Human MxB is localized in membraneless structures juxtaposed to cytoplasmic side of the nuclear pore (76). The antiviral activity of full-length MxB is evident against HIV and other lentiviruses which require RNA transit through the nuclear pore (74, 77). Steiner and Pavlovic report that MxB engineered to carry a nuclear localization signal is also antiviral towards FLUAV (74). In the case of mouse Mx proteins, the major MuMx1 is localized mainly in the nucleus, and we have previously shown that nuclear MuMx1 structures represent biomolecular condensates (46). It has been long been known that MuMx1 was antiviral against the nuclear-replicating FLUA, but not the cytoplasm-replicating VSV (40–42). Nevertheless, we discovered that a subset of Huh7 hepatoma cells (20-30% of transfected cells) showed that localization of MuMx1 along intermediate filaments in the cytoplasm. In such cells, we reported that MuMx1 was antiviral towards VSV (46).

An aspect of the significance of studies of MxA structure and antiviral function in response to hypoosmolar stress, especially in lung-derived cells, derives from the growing evidence of the protective role of IFN-α/β in covid-19 (23–31). An increase in MxA (often called “Mx1” in clinical papers) was observed in lung-derived tissues of patients with mild or moderate disease (29). Morbidity and mortality in this disease involves pulmonary edema and impaired ventilation, and is often observed in patients with defective IFN-α defense mechanisms (23, 30, 31). More generally, tissue-level edema is observed in hyponatremic hypoosmolar disorders such as salt-wasting nephropathies (e.g. polycystic kidney disease), mineralocorticoid deficiency (e.g. Addison’s disease), cirrhosis, and congestive heart disease which can comprise 15-20% of emergency room admissions (1–6). Such patients, with various underlying diseases, exhibit higher morbidity and mortality (2, 3, 5). The present study suggests that the IFN-α-induced MxA antiviral defense mechanism likely remains functional in the face of hypoosmolar stress.

## Experimental procedures

### Cells and cell culture

Human hepatoma cell line Huh7 was a gift from Dr. Charles M. Rice, The Rockefeller University. Human Hep3B, and A549 cells were obtained from the ATCC (Manassus, VA). Additional aliquots of the A549 cells and its derivative line A549-hACE2 was obtained from BEI Resources/ATCC (Manassus, VA). The respective cell lines were grown in DMEM (Corning Cat. No. 10-013-CV, with glutamine, Na-pyruvate and high glucose) supplemented with 10% v/v fetal bovine serum (FBS) in T25 flasks (44, 46). For experiments, the cells were typically plated in 35 mm dishes without or with cover-slip bottoms (44, 46). Recombinant human IFN-α2 was purchased from BioVision (Milpitas, CA) and typically used at 10-20 ng/ml for 2 days. Whole-cell extract and Western blot analyses for MxA were carried out as previously reported (43, 44).

### Plasmids and transient transfection

The GFP (1–248)-tagged full-length human MxA was a gift from Dr. Jovan Pavlovic (University of Zurich, Switzerland) (78); the expression vector for the HA-tagged full-length human MxA was a gift from Dr. Otto Haller (University of Freiburg, Germany) (79).; the HA and GFP tags were located on the N-terminal side of the Mx coding sequence. The expression vector for soluble RFP was a gift from Dr. Jason Lee (Baylor University School of Medicine, Houston, TX). Codon-optimized expression vactors for HA-tagged and GFP-tagged nucleocapsid (N) protein of SARS-CoV-2 virus were purchased from Sino Biologicals US Inc, Wayne, PA). Transient transfections were carried out using just subconfluent cultures in 35 mm plates using DNA in the range of 0.3-2 µg/culture and the Polyfect reagent (Qiagen, Germantown, MD) and the manufacturer’s protocol (with 10 µl Polyfect reagent per 35 mm plate).

### VSV stock and virus infection

A stock of the wild-type Orsay strain of VSV (titer: 9 x 10^8^ pfu/ml) was a gift of Dr. Douglas S. Lyles (Dept. of Biochemistry, Wake Forest School of Medicine, Winston-Salem, NC). Single-cycle virus infection studies at high multiplicity (moi >10 pfu/ml) were carried out essentially as described by Carey et al (67) as summarized in Davis et al (44) and in Sehgal et al (46). Briefly, A549 cultures (approx. 2 x 10^5^ cells per 35 mm plate), previously transfected with the pGFP-MxA expression vector (1-2 days earlier), were replenished with 0.25 ml serum-free Eagle’s medium and 10-20 µl of the concentrated VSV stock added (corresponding to MOI >10 pfu/cell). The plates were rocked every 15 min for 45 or 60 as indicated in respective experiments followed by addition of 2.5 ml of full culture medium. For the experiment shown in Suppl. Fig. 3, the cultures were subjected to a 5 min cycle of GFP-MxA condensate disassembly in ELB (verified by live-cell microscopy) and then returned to full culture medium till 4 hr post-infection (pi). For the experiment in Fig. 9, cultures infected for 45 min with VSV in isotonic DMEM, were shifted to full culture medium or one-third strength culture medium for another 4.25 hr. Cultures were fizxed in 4% PFA in isotonic PBS or one-third strength PBS (1 hr at 4°C immunostained for VSV nucleocapsid (N) protein using an mAb provided by Dr. Douglas S. Lyles (mAb 10G4). N-protein immunofluorescence (in red) in GFP-positive (nuclear or cytoplasmic) and negative cells was quantitated on a per cell basis as summarized in Davis et al (44) and Sehgal et al (46) and expressed in arbitrary units (AU) as integrated intensity/cell.

### Live-cell fluorescence imaging

Live-cell imaging of GFP-MxA structures in transiently transfected cells was carried out in cells grown in 35 mm plates using the upright the Zeiss AxioImager 2 equipped with a warm (37°C) stage and a 40x water immersion objective, and also by placing a coverslip on the sheet of live cells and imaging using the 100x oil objective (as above) with data capture in a time-lapse or z-stack mode (using Axiovision 4.8.1 software)(44).

### Phase transition experiments

Live GFP-MxA expressing cells in 35 mm plates were imaged using a 40x water-immersion objective 2-3 days after transient transfection in growth medium or serum-free DMEM medium or in phosphate-buffered saline (PBS). After collecting baseline images of Mx condensates (including time-lapse sequences), the cultures were exposed to 1,6-hexanediol (5% w/v) in PBS, or to hypotonic buffer (ELB; 10 mM NaCl, 10 mM Tris, pH 7.4, 3 mM MgCl_2_) and live-cell time-lapse imaging continued (44). After approximately 5-10 min the cultures exposed to hypotonic ELB were replenished with isotonic phosphate-buffered saline (PBS) and imaged for another 5-10 min. Time lapse images were also collected upon exposure of the cell cultures to different hypotonic buffers and treatments. Time lapse movies shown in Suppl. Movie 1 and 2 were imaged at one frame/5 sec.

### Immunofluorescence imaging

Typically, the cultures were fixed using cold paraformaldehyde (4%) for 1 hr and then permeabilized using a buffer containing digitonin or saponin (50 µg/ml) and sucrose (0.3M) (44). Single-label and double-label immunofluorescence assays were carried out using antibodies as indicated, with the double-label assays performed sequentially. Fluorescence was imaged as previously reported (44, 46) using an erect Zeiss AxioImager M2 motorized microscopy system with Zeiss W N-Achroplan 40X/NA0.75 water immersion or Zeiss EC Plan-Neofluor 100X/NA1.3 oil objectives equipped with an high-resolution RGB HRc AxioCam camera and AxioVision 4.8.1 software in a 1388 x 1040 pixel high speed color capture mode. Colocalization analyses were carried out using Image J software (Fiji) expressed in terms of Pearson’s correlation coefficient R.

### Quantitation of relative amounts of MxA in condensates vs dispersed state in a cell

The procedure used to quantitate the relative amounts of MxA in the condensed vs dispersed state on a per cell basis, as explained in Fig. 2 of ref. 38, is illustrated in Fig. 7B. Briefly, images with mixed condensate and dispersed MxA were subjected to Fourier filter processing to subtract objects of small radii (2-4 pixels) in Image J. The pixel radius (in the range 2-4 pixels) used for the subtraction was optimized to subtract all condensates from the image. MxA intensity in the residual subtracted image corresponded to the dispersed protein; subtracting this from the total intensity per cell gave the % of MxA in condensates on a per cell basis (see Fig. 7B).

### Antibody reagents

Rabbit pAb to human MxA (H-285) (ab-95926) was purchased from Abcam Inc. (Cambridge, MA); Mouse mAb to the VSV nucleocapsid (N) designated 10G4 was a gift from Dr. Douglas S. Lyles (Wake Forest School of Medicine, NC). Rabbit mAb to glyceraldehyde-3-phosphate dehydrogenase (GAPDH; 14C10; number 2118) was obtained from Cell Signaling (Danvers, MA), Anti-HA murine mAb (Cat. No. 100028-MM10) was purchased from Sino Biologicals US Inc. (Wayne, PA). Respective AlexaFluor 488- and AlexaFluor 594-tagged secondary donkey antibodies to rabbit (A-11008 and A-11012) or mouse (A-21202 and A-21203) IgG were from Invitrogen Molecular Probes (Eugene, OR). Alexafluor-594 tagged recombinant cholera toxin subunit B (CTB-red) was also purchased from Invitrogen Molecular Probes and used as per manufacturer’s recommendations (10 µM final concentration).

### Statistical testing

This was carried out (as in Fig. 9B and S3) using non-parametric one-way ANOVA (Kruskal-Wallis test) with Dunn’s post-hoc test for multiple comparisons.

## Abbreviations

2-DG: 2-deoxyglucose
CLEM: correlated live-cell fluorescence and electron microscopy
ELB: hypotonic “erythrocyte” lysis buffer
EM: thin-section
EM; FLUAV: influenza A virus
FRAP: fluorescence recovery after photobleaching assay
GAPDH: glycerandehyde 3-phosphate dehydrogenase
LLPS: liquid-liquid phase separation
MLO: membraneless organelle
Mx: myxovirus resistance protein
MuMx1: murine Mx1
MxA or HuMxa: human MxA
N protein: nucleocapsid protein
P bodies: cytoplasmic processing bodies
pi: post-infection
PM: plasma membrane
TEA: tetraethylammonium
TGT-020: *N*-1,3,4-thiadiazol-2-yl-3-pyridinecarboxamide
VLD: vacuole-like dilation
VSV: vesicular stomatitis virus

## Acknowledgements

We thank Drs. Joseph D. Etlinger (New York Medical College), Kenneth M. Lerea (New York Medical College) and Douglas S. Lyles (Wake Forest School of Medicine) for numerous helpful discussions. This research was supported, in part, by funds from New York Medical College, and by private funds of PBS.

## Author contributions

PBS designed the study, PBS carried out most of the experiments and imaging, HY carried out all the plasmid DNA preparations, HY and YJ carried out Western blotting, image quantitation and analyses, PBS wrote the manuscript. All authors approved the manuscript.

## Legends to Supplemental Data

**Supplemental Movie 1. Spontaneous reassembly of GFP-MxA condensates in an A549-hACE2 lung-cancer cell in one-fourth strength culture medium.** Movie shows time-lapse imaging (5 sec/frame) of an A549 cell expressing GFP-MxA condensates (three days after transfection) was commenced upon changing to one-fourth strength hypotonic culture medium.

**Supplemental Movie 2. Spontaneous reassembly of GFP-MxA condensates in an Huh7 hepatoma cell in one-fourth strength culture medium.** Movie shows time-lapse imaging (5 sec/frame) of an Huh7 expressing GFP-MxA condensates (two days after transfection) was commenced upon changing to one-fourth strength hypotonic culture medium.

**Supplemental Fig. 1.**
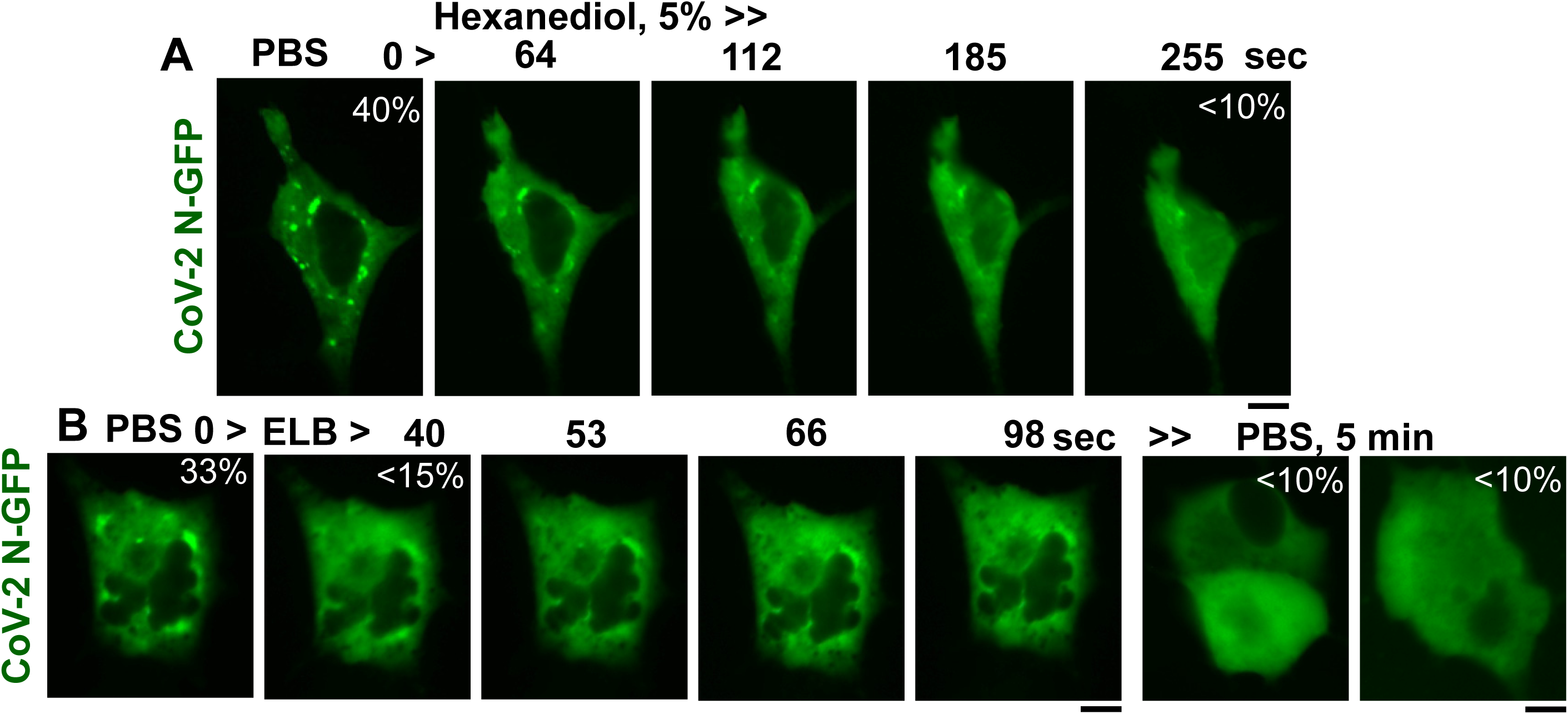
Properties of condensates formed by SARS-CoV-2 N-GFP. A549 cells in 35 mm plates were transiently transfected with an expression vector for SARS-CoV-2 N-GFP (Sino Biologicals) and the formation and properties of N-GFP condensates evaluated 2 days later. Panel A, Live-cell time-lapse images of condensates in a cell in isotonic PBS, and after shifting to 5% hexanediol in PBS. Panel B, Live-cell time-lapse images of condensates in a cell in isotonic PBS, and after shifting to hypotonic ELB; N-GFP-expressing cells in the same culture were also imaged 5 min after a further shift to isotonic PBS. All scale bars = 10 µm.

**Supplemental Fig. 2.**
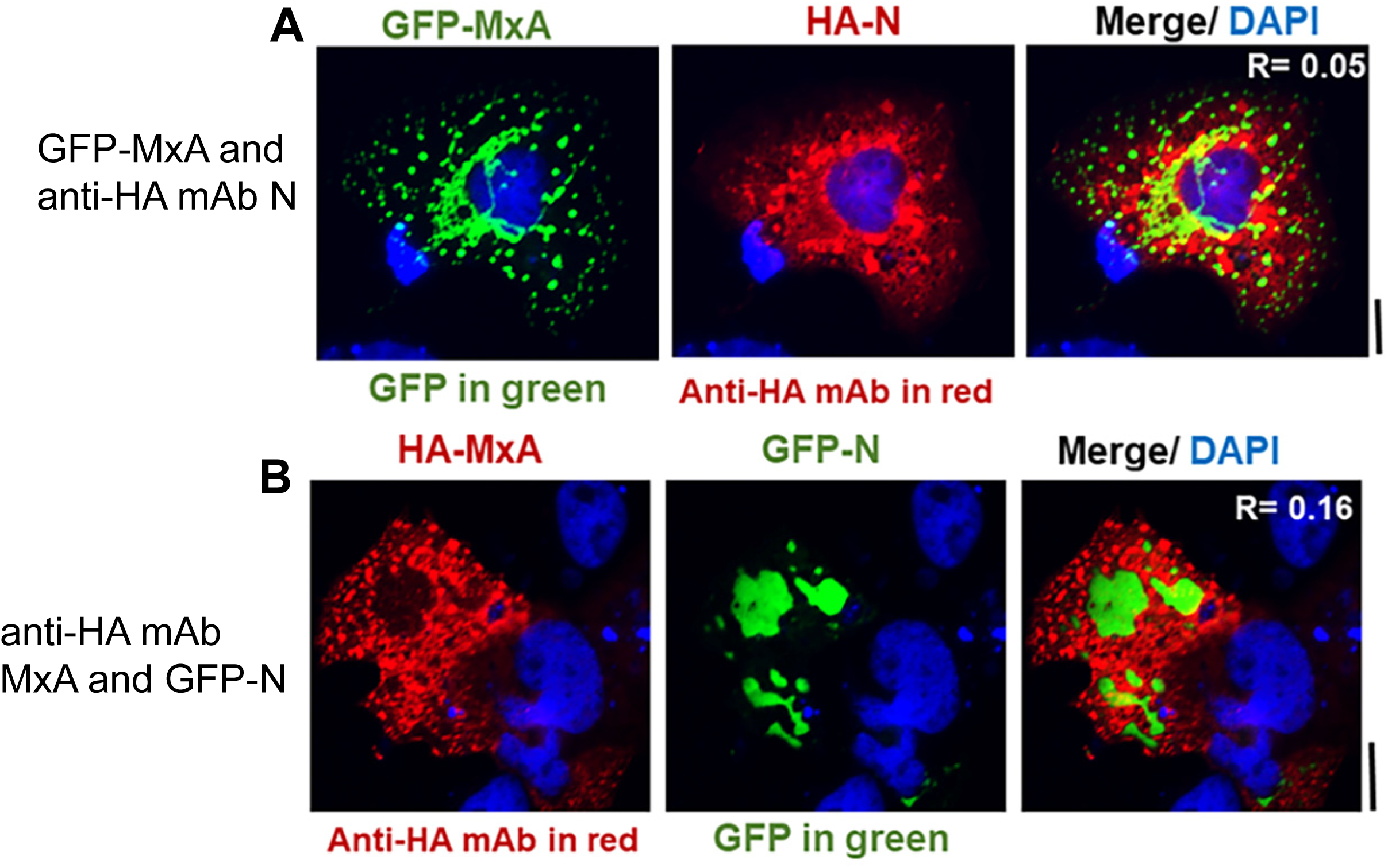
MxA and SARS-CoV-2 N protein form distinct condensates in the same cell. Huh7 cells in 35 mm plates were transiently transfected with pairwise combinations of respective GFP-tagged or HA-tagged vectors for MxA or SARS-CoV-2 N protein as indicated. The cultures were fixed (4% PFA) two days later, immunostained for HA (in red), and condensate formation by MxA and N protein evaluated by fluorescence imaging in green (for GFP) and red (for HA). R is the Pearson’s correlation coefficient between red and green; all scale bars = 10 µm.

**Supplemental Fig. 3.**
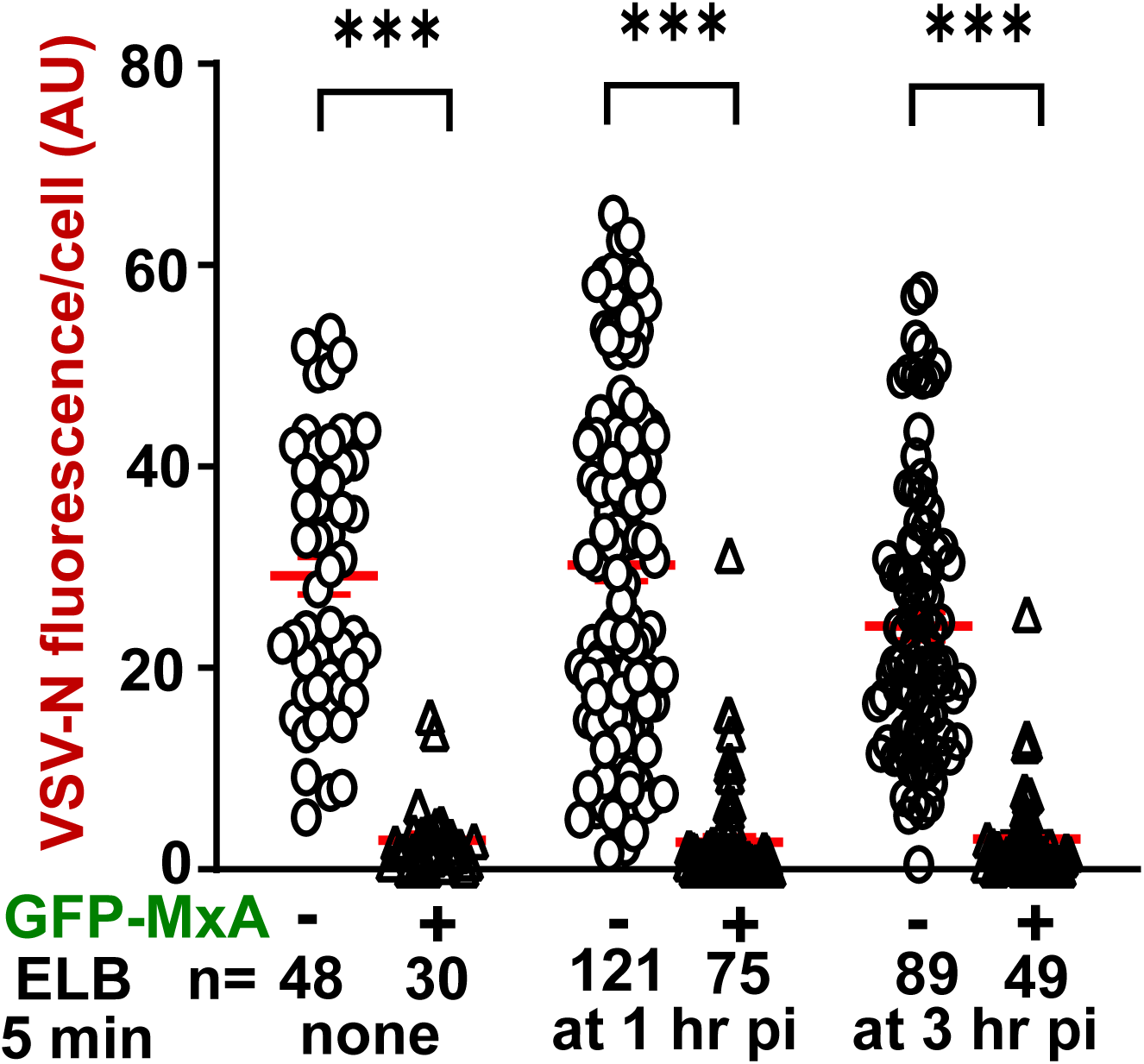
Antiviral activity of GFP-MxA against VSV in cultures subjected to one 5 min cycle of hypotonic disassembly/reassembly. Huh7 cells in 35 mm plates were transfected with the GFP-MxA vector, and 1 day later were infected with VSV (moi >10 pfu/cell)(44, 46). At one and 3 hr after the beginning of the virus infection (“post-infection”, indicated as pi in figure), respective cultures were shifted to hypotonic ELB for 5 min and then back to full medium. Live-cell microscopy confirmed the disassembly of GFP-MxA condensates when cell were shifted to ELB (not shown). All cultures were fixed at 4 hr pi, and immunostained for VSV N protein (in red)(as in refs. 44, 46). The VSV N protein fluorescence was quantitated in individual cells within the same culture without and with GFP-MxA (using Image J). n = number of cells evaluated per group in this experiment; horizontal red lines within each group indicate Mean ±SE. Statistical significance was evaluated using one-way ANOVA (Kruskal-Wallis with Dunn’s post-hoc test for multiple comparisons); *** *P* <0.001.

## References

1. Spasovski, G., Vanholder, R., Allolio, B., Annane, D., Ball, S., Bichet, D. Decaus, G., et al (2014) Clinical practice guidelines on diagnosis and treatment of hyponatremia. Eur J Endocrinology 170, G1–G47.

2. Walkar, S. S., Mount, D. B., Curhan, G. C. (2009) Mortality after hospitalization with mild, moderate, and severe hyponatremia. Am J Med 122, 857–865.

3. Verbalis, J. G. (2014) Hyponatremia and hypoosmolar disorders. National Kidney Foundation Primer on Kidney Diseases (sixth Edn), Elsevier, pgs. 62–70 doi: 10.1016/B978-1-4557-4617-0.00007-8

4. Rizzuto, W., Shemery, N., Bukowski, J. (2021) Acute water intoxication with resultant hypo-osmolar hyponatremia complicated by hypotension secondary to diffuse third spacing. Cureus 13, e18410 doi: 10.7759/cureus.18410

5. Adrogué, H. J., Madias, N. E. (2000) Hyponatremia. New Engl J Med 342, 1581–1589.

6. Danziger, J., Zeidel, M. L. (2015) Osmotic homeostasis. Clin J Am Soc Nephrol 10, 852–862.

7. Reuzeau, C., Mills, L. R., Harris, J. A., Morris, C. E. (1995) Discrete and reversible vacuole-like dilations induced by osmomechanical perturbations of neurons. J Membr Biol 145, 33–47.

8. Sheetz, M. P., Dai, J. (1996) Modulation of membrane dynamics and cell motility by membrane tension. Trends Cell Biol 6, 85–89.

9. Herring, T. L., Juranka, P., Mcnally, J., Lesiuk, H., Morris, C. E. (2000) The spectrin skeleton of newly-invaginated plasma membrane. J Muscle res Cell Motil 21, 67–77.

10. Barfod, E. T., Moore, A. L., Van de Graaf, B. G., Lidofsky, S. D. (2011) Myosin light chain kinase and Src control membrane dynamics in volume recovery from cell swelling. Mol Biol Cell 22, 634–651.

11. Gauthier, N. C., Fardin, M. A., Roca-Cusachs, P., Sheetz, M. P. (2011) Temporary increase in plasma membrane tension coordinates the activation of exocytosis and contraction during cell spreading. Proc Natl Acad Sci USA 108, 14467–14472.

12. Gauthier, N. C., Masters, T. A., Sheetz, M. P. (2012) Mechanical feedback between membrane tension and dynamics. Trends Cell Biol 22, 527–535.

13. Koffer, A., Williams, M., Johansen, T. (2002) Vacuole formation in mast cells responding to osmotic stress and to F-actin assembly. Cell Biol Internatl 26, 885–892.

14. Akella, R., Humphreys J. M., Sekulski K., He, H., Durbacz, M. et al (2021) Osmosensing by WNK kinases. Mol Biol Cell 32; 1614–1623.

15. Jalihal, A. P., Pitchiaya, S. Xiao, L., Bawa, m P., Jiang, X., Bedi, K., Parolia, A., Cieslik, M., Ljungman, M., Chinnaiyan, A. M., Walter, N. G. (2020) Multivalent proteins rapidly and reversibly phase-separate upon osmotic cell volume change. Mol Cell 79, 978–990.

16. Majumdar, S., Jain, A. (2020) Osmotic stress triggers phase separation. Mol Cell 79, 876–877.

17. Jalihal, A. P., Schmidt, A., Gao, G., Little, S. R., Pitchiaya, S., Walter, N. G. (2021) Hyperosmotic phase separation: condensates beyond inclusion, granules and organelles. J Biol Chem 296, 100044.

18. Knipe DM. 1990. Virus-host-cell interactions, p 293-316. *In* Fields BN, Knipe DM (ed), Fields’ Virology, 2nd ed, vol 1. Raven Press.

19. Fernandez-Puentes C, Carrasco L. 1980. Viral infection permeabilizes mammalian cells to protein toxins. Cell 20:769–75.

20. Carrasco L. 1978. Membrane leakiness after viral infection and a new approach to the development of antiviral agents. Nature 272:694–9.

21. Carrasco L, Smith AE. 1976. Sodium ions and the shut-off of host cell protein synthesis by picornaviruses. Nature 264:807–9.

22. Montasir M, Rabin ER, Phillips CA. 1966. Vaccinia pneumonia in mice. A light and electron microscopic and viral assay study. Am J Pathol 48:877–95.

23. Decker, T. (2021) The early interferon catches the SARS-CoV-2. J Exp Med 218, e20211667

24. Cheemarla, N., Watkins, T. A., Mihaylova, V. T., Wang, B., Zhao, D., Wang, G., Landry, M. L., Foxman, E. F. (2021) Dynamic innate immune response determines susceptibility to SARS-CoV-2 infection and early replication kinetics. J Exp Med 218, e20210583

25. Rarani, F. Z,, Rarani, M. Z., Hamblin, M. R., Rashidi, B., Hashemian, S. M. R., Mirzaei, H. (2022) Comprehensive overview of COVID-19-related respiratory failure: focus on cellular interactions. Cell Mol Biol Lett 27, 63. Doi: 10.1186/s11658-022-00363-3

26. Mantlo, E., Bukreyeva, N., Maruyama, J., Paessler, S. and Huang, C. (2020) Antiviral activities of type I interferons to SARS-CoV-2 infections. Antiviral Research 179, 104811.

27. Andreakos, E. and Tsiodras, S. (2020) Covid-19: lambda interferon against viral load and hyperinflammation, in press. doi: 10.15252/emmm.202012465

28. Felgenhauer, U., Schoen, A., Gad, H. H., Hartmann, R., Schaubmar, A. R., Failing, K., Drosten, C., Weber, F. (2020) Inhibition of SARS-CoV-2 by type I and type III interferons. J Biol Chem 295, 13958–13964.

29. Bizzotto, J, Sanchis, P., Abbate, M., Lage-Vickers, S., Lavignolle, R., et al SARS-CoV-2 infection boosts *MX1* antiviral effector in COVID-19 patients. iScience23, 101585

30. Kwon, D. (2021) Rogue antibodies linked to deaths in from severe COVID. Nature 597, 162

31. Bastard, P., Orlova, E., Sozaeva, L., Levy R., James, A. et al (2021) Preexisting autoantibodies to type I IFNs underlie critical COVID-19 pneumonia in patients with ARA-1. J Exp Med 218, e20210554

32. Mitrea, D.M. and Kriwacki, R.W. (2016) Phase separation in biology, functional organization of a higher order. Cell Commun. Signal 14, 1. doi:10.1186/s12964-015-0125-7.

33. Banani, S.F., Lee, H.O., Hyman, A.A. and Rosen, M.K. (2017) Biomolecular condensates: organizers of cellular biochemistry. Nat Rev Mol Cell Biol. 18, 285–298.

34. Shin, Y. and Brangwynne, C.P. (2017) Liquid phase condensation in cell physiology and disease. Science 357(6357) pii: eaaf4382. doi: 10.1126/science.aaf4382.

35. Alberti, S. (2017) The wisdom of crowds: regulating cell function through condensed states of living matter. J Cell Sci. 130, 2789–2796.

36. Gomes, E. and Shorter. J. (2019) The molecular language of membraneless organelles. J Biol Chem 294, 7115–7127.

37. Alberti, S., Gladfelter, A. and Mittag, T. (2019) Considerations and challenges in studying liquid-liquid phase separation and biomolecular condensates. Cell 176, 419–434.

38. Sehgal, P. B., Westley, J., Lerea, K. M. DiSenso-Browne, S. and Etlinger, J. D. (2020) Biomolecular condensates in cell biology and virology: phase-separated membraneless organelles (MLOs). Analytical Biochem 597, 113691.

39. Sehgal, P. B. (2021) Metastable biomolecular condensates of the interferon-inducible antiviral Mx-family GTPases: a paradigm shift in the last three years. J Biosci 46, 72 doi: 10.1007/s12038-021-00187-x

40. Haller, O. and Kochs, G. (2002). Interferon-induced mx proteins: dynamin-like GTPases with antiviral activity. Traffic 3, 710–717.

41. Haller, O., Staeheli, P., Kochs, G. (2007) Interferon-induced Mx proteins in antiviral host defense. Biochimie 89, 812–818.

42. Haller, O., Staeheli, P., Schwemmle, M. and Kochs, G. (2015) Mx GTPases: dynamin-like antiviral machines of innate immunity. Trends Microbiol 23, 154–163.

43. Davis, D., Yuan, H., Yang, Y.M., Liang, F.X. and Sehgal. P.B. (2018) Interferon-alpha-induced cytoplasmic MxA structures in hepatoma Huh7 and primary endothelial cells. Contemp Oncol (Pozn) 22, 86–94.

44. Davis, D., Yuan, H., Liang, F.X., Yang, Y.M., Westley, J., Petzold, C., Dancel-Manning, K., Deng, Y., Sall, J. and Sehgal, P.B. (2019) Human antiviral protein MxA forms novel metastable membraneless cytoplasmic condensates exhibiting rapid reversible tonicity-driven phase transitions. J Virol 93, e01014–19.

45. Kochs, G., Janzen, C., Hohenberg, H. and Haller, O. (2002) Antivirally active MxA protein sequesters La Crosse virus nucleocapsid protein into perinuclear complexes. Proc Natl Acad Sci USA 99, 3153–3158.

46. Sehgal, P. B., Yuan, H., Scott, M. F., Deng, Y., Liang, F-X., Mackiewicz, A (2020) Murine GFP-Mx1 forms nuclear condensates and associates with cytoplasmic intermediate filaments: novel antiviral activity against VSV. J Biol Chem 295, 18023–18035.

47. Ndubuisi, M. I., Guo, G. G., Fried, V. A., Etlinger, J. D., Sehgal, P. B. (1999) Cellular physiology of STAT3: where’s the cytoplasmic monomer? J Biol Chem 274, 25499–25509.

48. Sehgal, P. B. (2019) Biomolecular condensates in cancer cell biology: interleukin-6-induced cytoplasmic and nuclear STAT3/PY-STAT3 condensates in hepatoma cells. Contemp Oncol (Pozn). 23, 16–22.

49. Kochs, G., Haener, M., Aebi, U., Haller, O. (2002) Self-assembly of human MxA GTPase into highly ordered dynamin-like oligomers. J Biol Chem 277, 14172–14176.

50. Haller, O., Gao, S., von der Malsburg, A., Daumke, O., Kochs, G. (2010) Dynamin-like MxA GTPase: structural insights into oligomerization and implications for antiviral activity. J Biol Chem 285, 28419–28424,

51. Nigg, P. E., Pavlovic, J. (2015) Oligomerization and GTP-binding requirements of MxA for viral target recognition and antiviral activity against influenza A virus. J Biol Chem 290, 29893–29906.

52. Dick, A., Graf, L., Olal, D., von der Malsburg, A., Gao, S., Kochs, G., Daumke, O. (2015) Role of nucleotide binding and GTPase domain dimerization in dynamin-like myxovirus resistance protein A for GTPase activation and antiviral activity. J Biol Chem 290, 12779–12792.

53. Graf, L., Dick, A., Sendker, F., Barth, E., Marz, M., Daumke, O., Kochs, G. (2018) Effects of allelic variations in the human myxovirus resistance protein A on its antiviral activity. J Biol Chem 293, 3056–3072.

54. Jain, S., Wheeler, J. R., Walters, R. W., Agrawal, A., Barsic, A., Parker, R. (2016) ATPase-modulated stress granules contain a diverse proteome and substructure. Cell 164: 487–498.

55. Mateju, D., Franzmann, T. M., Patel, A., Kopach, A., Boczek, E. E., Maharana, S., Lee, H. O., Carra, S., Hyman, A. A., Alberti, S. (2017) An aberrant phase transition of stress granules triggered by misfolded protein and prevented by chaperone function. EMBO J 36: 1669–1687

56. Ditlev, J. A., Case, L. B. and Rosen, M. K. (2018) Who’s in and who’s out - compositional control of biomolecular condensates. J. Mol. Biol. 430: 4666–4684.

57. Lee, J., Reich, R., Xu, F., Sehgal, P. B. (2009) Golgi, trafficking, and mitosis dysfunctions in pulmonary arterial endothelial cells exposed to monocrotaline pyrrole and NO scavenging. Am J Physiol Cell Mol Physiol 297, L715–L728.

58. Singh, V., Xu, L., Boyko, S., Surewicz, K., Surewicz, W. K. (2020) Zinc promotes liquid-liquid phase separation of tau protein. J Biol Chem 295, 5850–5856.

59. Cubuk, J., Alston, J. J., Incicco, J. J., Singh, S., Stuchell-Brereton, M. D., Ward, M. D., Zimmerman, M. I., Vithani, N., Griffith, D., Wagoner, J. A., Bowman, G. R., Hall, K. B., Soranno, A. and Holehouse, A. S. (2021) The SARS-CoV-2 nucleocapsid protein is dynamic, disordered, and phase separates with RNA, Nature Commun 12, 1936

60. Perdikari, T. M., Murthy, A. C., Ryan, V. H., Watters, S., Naik, M. T. and Fawzi, N. L. (2020) SARSCoV-2 nucleocapsid protein undergoes liquid-liquid phase separation stimulated by RNA and partitions into phases of human ribonucleoproteins, EMBO J 39, e106478

61. Iserman, C., Roden, C., Boerneke, M., Sealfon, R., McLaughlin, G., Jungreis, I., Park, C., Boppana, A., Fritch, E., Hou, Y. J, Theesfeld, C., Troyanskaya, O. G., Baric, R. S., Sheahan, T. P., Weeks, K. and Gladfelter, A. S. (2020) Genomic RNA elements drive the phase separation of the SARS CoV-2 nucleocapsid *Mol Cell* 80: 1078-1091.e6

62. Carlson, C. R., J.B. Asfaha, C.M. Ghent, C.J. Howard, N. Hartooni, D.O. Morgan (2020) Phosphoregulation of phase separation by the SARS-CoV-2 N protein suggests a biophysical basis for its dual functions. Mol Cell 80: 1092–1103.e4

63. Lausel, C.’ Leon, S. (2020) Cellular toxicity of the metabolic inhibitor 2-deoxyglucose and associated resistance mechanisms. Biochem Pharmacol 182, 114213.

64. Rai, A. K., Chen, J-X., Selbach, M., Pelkmans, L. (2018) Kinase-controlled phase transition of membraneless organelles in mitosis. Nature 559, 211–216.

65. Risso-Ballester, J., Galloux, M., Cao, J., La Groffic, R., Hontonnou, F., Jobart-Malfait, A. et al. (2020) A condensate-hardening drug blocks RSV replication in vivo. Nature 595, 596–599.

66. Wagner, K., Unger, L., Salman, M. M., Kitchen, P., Bill, R. M., Yool, A. J. (2022) Signaling mechanisms and pharmacological modulators governing diverse aquaporin functions in human health and disease. Int J Med Sci 23, 1388 doi: 10.3390/ijms23031388

67. Carey BL, Ahmed M, Puckett S, Lyles DS. 2008. Early steps of the virus replication cycle are inhibited in prostate cancer cells resistant to oncolytic vesicular stomatitis virus. J Virol 82:12104–12115.

68. Holzwarth, G., Bhandari, A., Tommervik, L., Macosko, J. C., Ornellas, D. A., Lyles, D. S. (2020) Vesicular stomatitis virus nucleocapsids diffuse through cytoplasm by hopping from trap to trap in random directions. SciReports 10: 10643.

69. Yuan, H., Sehgal, P. B. (2016) MxA is a novel regulator of endosome-associated transcriptional signaling by bone morphogenetic proteins 4 and 9 (BMP4 and BMP9). PLoS ONE 11, e0166382

70. Kar, M., Dar, F., Welsh, T. J., Vogel, L. T., Kuhnmuth, R., Majumdar, A., Krainer, G. et al (2022) Phase-separating RNA-binding proteins form heterogeneous distributions of clusters in subsaturated solutions. Proc Natl Acad Sci USA 119, e2202222119

71. Fuxreiter, M. (2022) Electrostatisc tunes protein interactions to context. Proc Natl Acad Sci USA 119, e2209201119

72. Watanabe, K., Saito, K., Kinjo, M., Matsuda, T., Tamura, M., Kon, S., Miyazaki, T., Uede, T. (2004) Molecular dynamics of STAT3 on IL-6 signaling pathway in living cells. Biochem Biophys Res Commun 324, 1264–1273.

73. Lussier, M. P., Cayoutte, S., Lepage P. K., Bernier, C. L., Francoer, N., St. Hilare, M., Pinard, M., Boulay, G. (2005) MxA, a member of the dynamin superfamily, interacts with the ankyrin-like repeat domain of TRPC. J Biol Chem 280, 19393–19400.

74. Steiner, F., Pavlovic, J. (2020) Subcellular localization of MxB determines its antiviral potential against influenza virus. J Virol 94, e00125–e220

75. Zürcher, T., Pavlovic, J. and Staeheli, P. (1992) Mechanism of human MxA protein action: variants with changed antiviral properties. EMBO J 11: 1657–1661.

76. King, M. C., Raposo, G. and Lemmon, M. A. (2004) Inhibition of nuclear import and cell-cycle progression by mutated forms of the dynamin-like GTPase MxB. Proc. Natl. Acad. Sci. USA 101: 8957–8962.

77. Liu, Z., Pan, Q., Ding, S., Qian, J., Xu, F., Zhou, J., Cen, S., Guo, F. and Liang, C. (2013) The interferon-inducible MxB protein inhibits HIV-1 infection. Cell Host & Microbe 14: 398–410.

78. Wisskirchen C, Ludersdorfer TH, Muller DA, Moritz E, Pavlovic J. 2011. Interferon-induced antiviral protein MxA interacts with the cellular RNA helicases UAP56 and URH49. J Biol Chem 286:34743–34751.

79. Stertz, S., Reichelt, M., Krijnse-Locker, J., Mackenzie, J., Simpson, J. C., Haller, O. and Kochs, G. (2006). Interferon-induced, antiviral human MxA protein localizes to a distinct subcompartment of the smooth endoplasmic reticulum. J. Interferon Cytokine Res. 26(9):650–660.

